# Half-life of biodegradable plastics in the marine environment depends on material, habitat, and climate zone

**DOI:** 10.1101/2021.01.31.429013

**Authors:** Christian Lott, Andreas Eich, Dorothée Makarow, Boris Unger, Miriam van Eekert, Els Schuman, Marco Segre Reinach, Markus T. Lasut, Miriam Weber

**Affiliations:** HYDRA Marine Sciences GmbH, Bühl, Germany; HYDRA Fieldwork, Sulzburg, Germany; LeAF BV, Wageningen, The Netherlands; Coral Eye Outpost, Pulau Bangka, Sulawesi Utara, Indonesia; Sam Ratulangi University UNSRAT, Manado, Sulawesi Utara, Indonesia

**Keywords:** Mediterranean Sea, Southeast Asia, surface erosion rate, environmental persistence, lifecycle assessment, polyhydroxyalkanoate, polybutylene sebacate, polybutylene sebacate co-terephthalate

## Abstract

The performance of the biodegradable plastic materials polyhydroxyalkanoate (PHA), polybutylene sebacate (PBSe) and polybutylene sebacate co-terephthalate (PBSeT), and of polyethylene (LDPE) was assessed under marine environmental conditions in a three-tier approach. Biodegradation lab tests (20 °C) were complemented by mesocosm tests (20 °C) with natural sand and seawater and by field tests in the warm-temperate Mediterranean Sea (12 – 30 °C) and in tropical Southeast Asia (29 °C) in three typical coastal scenarios. Plastic film samples were exposed in the eulittoral beach, the pelagic open water and the benthic seafloor and their disintegration monitored over time. We used statistical modelling to predict the half-life for each of the materials under the different environmental conditions to render the experimental results numerically comparable across all experimental conditions applied. The biodegradation performance of the materials differed by orders of magnitude depending on climate, habitat and material and revealed the inaccuracy to generically term a material ‘marine biodegradable’. The half-life *t*_0.5_ of a film of PHA with 85 μm thickness ranged from 54 d on the seafloor in SE Asia to 1247 d in mesocosm pelagic tests. *t*_0.5_ for PBSe (25 μm) ranged from 99 d in benthic SE Asia to 2614 d in mesocosm benthic tests, and for PBSeT *t*_0.5_ ranged from 147 d in the mesocosm eulittoral to 797 d in Mediterranean benthic field tests. For LDPE no biodegradation could be observed. These data can now be used to estimate the persistence of plastic objects should they end up in the marine environments considered here and will help to inform the life cycle (impact) assessment of plastics in the open environment.

## 1 Introduction

Global plastic production is growing exponentially. It has almost doubled since the beginning of this century (Plastics Europe, 2019) to 400 Mt in 2020 (including fibers) and is estimated to reach 800 Mt in 2050 (Rouch, 2019). Biodegradable polymers are a small, but growing segment in this market with a share of 0.3 % (1.227 Mt) in 2020 (European Bioplastics, 2019). The amount of plastic entering the natural environment is augmenting rapidly, accumulating at an exponential rate (Geyer et al., 2017). If introduced to the environment, e.g. as litter, it can be assumed that most items made from biodegradable plastic materials have similar pathways and sinks as conventional non-biodegradable plastic items. Plastic pollution is found almost anywhere in nature it has been looked for, including air (e.g. Dris et al., 2016), high-mountain and polar ice (e.g. Ambrosini et al., 2019, Kanhai et al., 2020), terrestrial soil (e.g. review by Helmberger et al., 2019), freshwater (e.g. Wagner et al., 2014) and marine systems (e.g. Weber et al., 2015, fig. 2 and references therein) with effects on ecosystem level, organism level and on humans still to be fully understood.

Biodegradable plastics are used as alternative materials to conventional plastics, e.g. for agricultural films (e.g. Sintim and Flury, 2017) and/or as substitutes required by legislation such as fruit and vegetable bags in Italy and France (Gazzetta Ufficiale della Repubblica Italiana, 2017, Journal Officiel de la République Française, 2016). Biodegradable polymers are discussed as a mitigation strategy against environmental plastic pollution and e.g. the government of the Republic of Indonesia in the National Plan of Action on Marine Plastic Debris 2017-2025 called to increase the share of biodegradable plastics to fight plastic pollution of the ocean (Republic of Indonesia, 2017). The European Commission in their European Plastics Strategy (European Commission, 2018) stated that new plastics with biodegradable properties bring new opportunities but also risks. It acknowledged the role of biodegradable plastics in some applications and that studies are needed to develop a clear regulatory framework and a lifecycle assessment to identify the criteria for such applications (Albertsson et al., 2020). It was also pointed out the importance to make sure that biodegradable plastics are not put forward as a solution to littering.

For some applications where the unintentional loss of plastic to the environment is intrinsic to its use (e.g. fishing gear, boating gear, and beach tourism items) or which are prone to unavoidable input (e.g. abrasion of tires, shoes, textiles, paint) and thus are continuously introduced to the environment, biodegradable polymeric alternatives might be the only solution from the material side (Albertson et al., 2020).

There is no universal definition, yet several descriptions and definitions of the term ‘biodegradation’ exist, which might lead to decision-making based on wrong assumptions and even to misuse, false claims, and disinformation. Following the biogeochemical point of view, we define ‘biodegradation’ of a polymer as the mineralization to carbon dioxide (and in the absence of oxygen also methane), water, and the incorporation of its breakdown products into new biomass by naturally occurring bacteria, archaea, and fungi, leaving no residue behind.

Hence, ‘biodegradability’ describes the capacity of a polymeric material to be broken down by microorganisms in the considered receiving environment. As the abundance, diversity, and activity of microorganisms vary in nature as do the environmental conditions, also the specific biodegradation at a given place and time will vary. To be truly meaningful, the term ‘biodegradable’ must therefore be clarified and linked not only to a duration in time, compatible with a human scale but also to conditions of biodegradation, and to be seen as a system property (Albertsson et al., 2020).

As reliable, comparable, and verifiable information is needed and officially asked for (The European Green Deal, European Commission, 2019) the claim ‘biodegradable’ of a certain material should be sufficiently specified and reliably proven. Therefore suitable tests are needed.

Here, we tested the performance of biodegradable plastic in the marine environment applying laboratory methods proposed by Tosin et al. (2012) and field and mesocosm methods developed during the EU project Open-Bio (Lott et al., 2016a and 2016b, 2020). In a 3-tier approach, we answered the questions whether the tested materials are biodegradable at all, whether biodegradation does take place under real natural marine conditions, and at which rates in the different environmental settings.

To enable a numerical comparison of the experimental results we applied a statistical model to the experimental results to mathematically describe the biodegradation over time with a specific half-life. This number can further be used as a material property specific for defined environmental conditions and used to set thresholds, estimate environmental risk, and be fed into lifecycle assessment.

In this study, we tested three biodegradable polymers with natural marine seawater and sediments in lab and mesocosm tests, and in field tests in three coastal marine scenarios in the Mediterranean Sea and in tropical Southeast Asia. We focused on three easily accessible habitats in coastal shallow water: the intertidal beach, the open water, and the sandy seafloor.

## 2 Materials and methods

### 2.1 Test materials

Four polymer materials were selected to assess their biodegradation in the different test systems: polyhydroxyalkanoate copolymer, a bacteria-derived, thus bio-based, biodegradable material as a positive control, polybutylene sebacate (PBSe), polybutylene sebacate co-butylene terephthalate (PBSeT), two polymers commonly used in blends for plastic products, and low-density polyethylene (LDPE) as the negative control (Table 1).

**Table 1.** List of tested polymers with their properties, film thickness, compounds, and supplier. Percentage of total organic carbon (TOC), total carbon (TC), hydrogen (H), and nitrogen (N) analyzed with standard methods.

| Test material | Thickness, compounds, supplier | TOC (%) | TC (%) | H (%) | N (%) |
| --- | --- | --- | --- | --- | --- |
| Low Density Polyethylene<br>**LDPE**<br>(negative control) | Film 30 microns<br>Grade: LUPOLEN 2420K, LyondellBasell | 85.03 | 85.37 | 14.68 | < 0.1 |
| Polybutylene Sebacate<br>**PBSe** | Film 25 microns<br>aliphatic polyester | 65.26 | 65.58 | 7.69 | < 0.1 |
| Polybutylene Sebacate<br>co-butylene Terephthalate<br>**PBSeT** | Film 25 microns<br>aliphatic-aromatic polyester | 65.25 | 65.81 | 9.54 | < 0.1 |
| Polyhydroxyalkanoate Copolymer<br>**PHA**<br>(positive control) | Film 85 microns<br>Grade: Mirel™ P5001, Metabolix<br>compound >70% PHA copolymer,<br>plasticizer, fillers | 47.82 | 49.11 | 6.03 | 0.52 |

The materials were tested as films. In the lab tests, small pieces of film (4 × 4, 2 × 4,3 × 3, ø 2 cm) were directly incubated in the test vessels. For the mesocosm and field tests polymer film samples were mounted in HYDRA® test frames (260 × 200 mm external and 200 × 160 mm internal dimensions leaving a surface of 320 cm^2^ of material directly exposed), i.e. held between mesh (PET) and plastic frames (PE) (SI Figure 1) to prevent mechanical impact on the sample, as described before (Lott et al., 2020).

**Figure 1.**
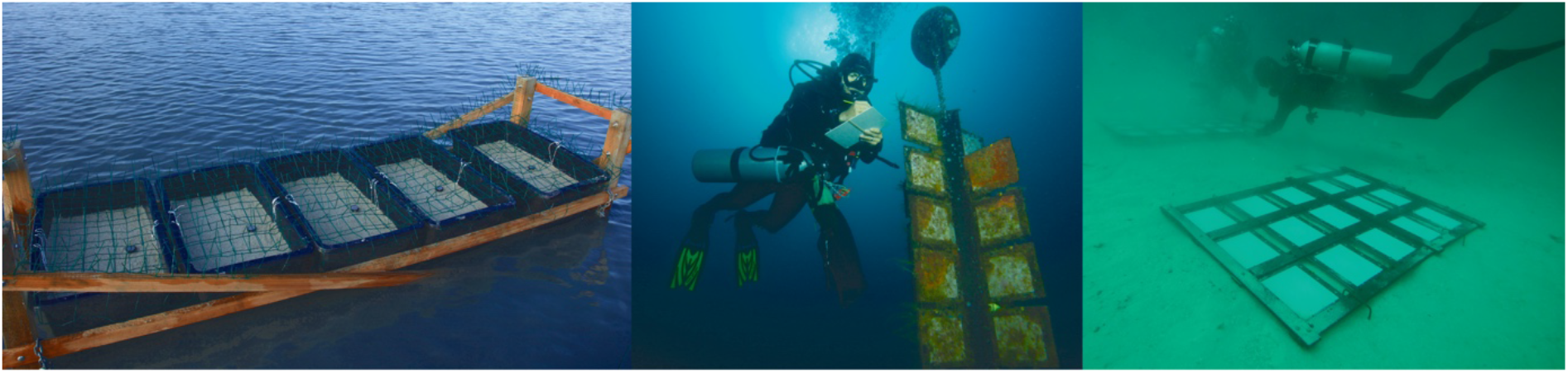
Field test systems: left: eulittoral test: Samples were exposed in bins filled with beach sand on a rack in the midwater line, center: pelagic tests: samples attached to a floating rack exposed at 20 m water depth, right: benthic tests: samples were mounted in panels that were fixed on the sandy seafloor at 40 m (Mediterranean Sea) and 32 m water depth (SE Asia)

**Figure 2.**
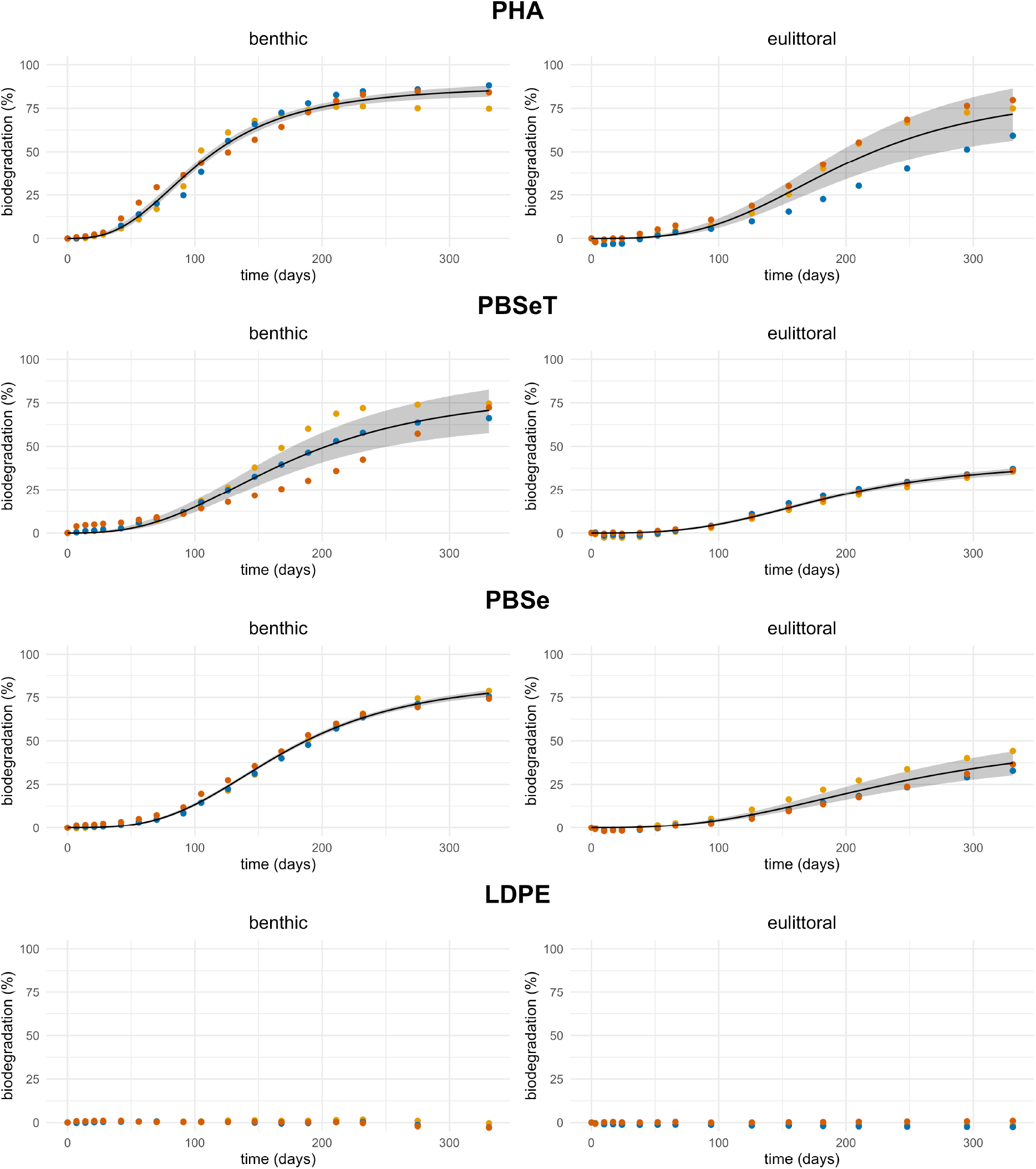
Laboratory experiments with Mediterranean Sea water and sediment: Biodegradation curves of PHA, PBSe, and PBSeT. Biodegradation of tested polymers of polyhydroxyalkanoate (PHA) as positive control, test materials polybutylene sebacate co-terephthalate (PBSeT) and polybutylene sebacate (PBSe) and low-density polyethylene (LDPE) as negative control when incubated with seafloor sediment at the sediment-seawater interface (benthic) and buried in beach sand (eulittoral). Colors red, yellow, and blue indicate samples from different test flasks, dots represent raw data analyzed in the model. The model curve ± 95% CI (shaded area) was drawn if estimated parameters were significantly different from 0.

### 2.2 Matrices sediment and seawater

Eulittoral sediment: Natural marine sediment for the eulittoral tests in the lab and in mesocosms was retrieved at about 0.1 m water depth from the beach of Fetovaia, Isola d’Elba, Italy, (42°44’00.1“N, 010°09’15.3“E) and is called ‘beach sediment’. Sublittoral sand: Carbonate sediment for the benthic mesocosm tests was collected by divers from the seafloor at 40 m depth off Isola di Pianosa, National Park Tuscan Archipelago, Italy, (42°34’41.4“N, 010°06’30.6“E) and is called ‘seafloor sediment’. Larger pieces, like plant material, sea shells, pieces of driftwood, etc. were removed by sieving through a 10 mm mesh after collection. Seawater was taken at Seccheto, Isola d’Elba, Italy (42°44’06.5“N, 010°10’33.5“E) and was used for the lab experiments, to wash the sediments and to fill the mesocosms. The mesocosm tests were run twice for about one year each and specified as y1 and y2 in this text. The matrices were renewed after the first run and the physical and chemical properties of sand and water were analyzed with standard methods at the beginning of each experiment (for details see Lott et al., 2020). The water used in the mesocosm tests (see below) had low to moderate levels of organic carbon, nitrogen and phosphor compounds, and chlorophyll. No toxic substances such as heavy metals, organotin compounds, or persistent organic pollutants (POPs) were detected. The beach sand used for tests was in the grain size range of medium sand and of mainly siliclastic origin. The seafloor sediment used for the benthic experiments was mainly carbonate fine sand. Porosity and permeability were slightly lower for both sand types in y2 (Table 2). Metal concentrations were low or below the detection limit. The test for persistent organic pollutants (POPs) in the sediments used for the experiments was negative (details in Lott et al., 2020).

**Table 2.**
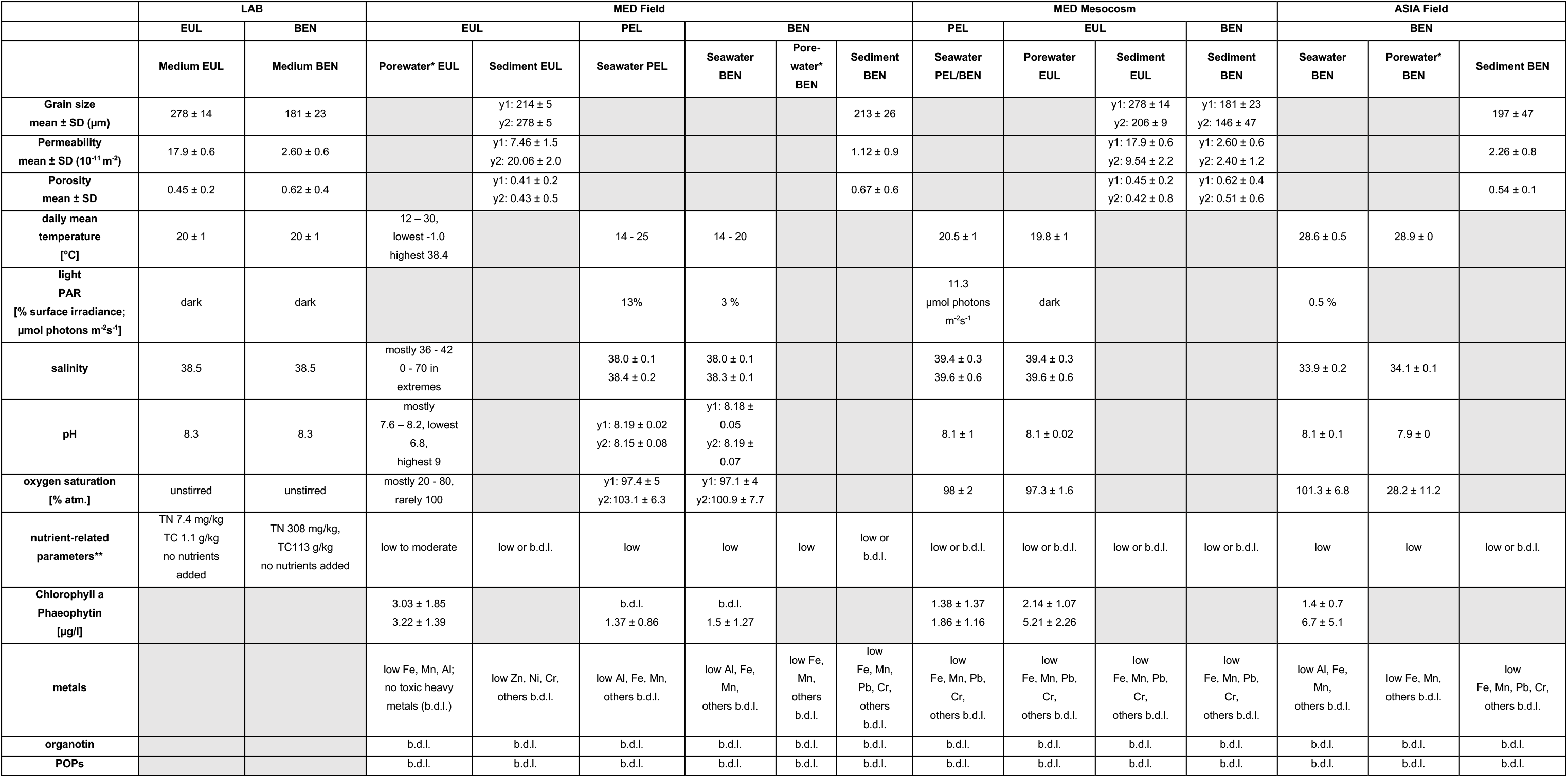
Experimental conditions: Properties of the water and sediments from the lab, field, and mesocosm experiments. *Porewater was taken at 5 cm sediment depth; **nutrient-related parameters *sensu* Weber et al., (2012), LAB = laboratory tests, MED = Mediterranean Sea, ASIA = tropical SE Asia, EUL = eulittoral (beach scenario), BEN = benthic (sublittoral sand bottom scenario), PEL = pelagic (open water scenario), low = values just above detection limit, moderate = values within one order of magnitude of detection limit, b.d.l. = below detection limit, grey shaded fields = not analyzed, TN = total nitrogen, TC = total carbon, PAR = photosynthetically active radiation (400 – 1000 nm), POPs = persistent organic pollutants.

### 2.3 Laboratory tests

The lab tests were conducted at LeAF in Wageningen, Netherlands, with water and sediment from the Mediterranean field test locations (Table 2). The matrices were collected in plastic containers, shipped to LeAF and stored at 4°C until further use. At the start of the experiments, water and sediment were characterized with standard analytical methods.

#### 2.3.1 Eulittoral test (beach scenario)

The plastic samples were buried in 400 g of beach sediment under aerobic conditions as described in Tosin et al. (2012), following ISO 22404 (2019). The sediment had a water content of 18.9%, total solids content of 81.1%, of which volatile solids of 0.57%, total nitrogen of 7.4 mg/kg, and total carbon of 1.1 g/kg. No nutrients were added. The tests were carried out in 2-L Duran® wide-mouth bottles and a container for the CO_2_ sorption connected to a side port. The container was filled with 30 mL of 0.5N KOH solution. The bottle and the container were closed with a Python rubber stopper (Rubber BV, Hilversum, The Netherlands). The bottles were incubated in the dark in a closed box that was placed in a temperature-controlled room (20 ± 1 °C). The test materials (PBSe and PBSeT) were square-shaped specimens with a dimension of approximately 40 × 40 mm. The negative control was done with 40 × 40 mm specimens of LDPE. For the test with PHA as the positive control specimens were cut in 20 × 40 mm pieces (because of the high grammage). The mass of each specimen (about 100 mg) was recorded. For the test 15 reactors were prepared to enable testing in triplicate of PBSe, PBSeT, PHA (reference material/positive control), LDPE (negative control), and blanks to correct for endogenous respiration. All reactors were pre-incubated without plastic samples to assess the endogenous respiration. After one week of pre-incubation, the CO_2_ production was measured. The results showed that the endogenous respiration was similar in the bottles. After the pre-incubation, the test was started. For this, the reactors were opened and 100 g of sediment was removed. The sediment surface was smoothened and two specimens of test materials PBSe or PBSeT or LDPE or one specimen of PHA was placed on the surface. Thereafter the withdrawn sediment was carefully put on top of the sediment and test material. The specimens were covered with sand in a homogenous layer. The blanks for endogenous respiration did not receive any test specimen.

The extent of biodegradation (i.e. (endogenous) respiration) was assessed by determining the amount of carbon dioxide produced and absorbed in the KOH solution by titration with 0.3N HCl solution to pH 8 and thereafter further to pH 3.8. The amount of CO_2_ absorbed was calculated according to the formula given by Tosin et al. (2012). The container for the CO_2_ absorber was removed and analyzed and titrated before its capacity was exceeded. Each time the KOH was replaced by a fresh solution the reactor was weighed to monitor moisture loss from the sediment and allowed to sit open for approximately 15 min so that the air in the reactor was refreshed before resealing the reactor. Distilled water was added back periodically to the sediment to maintain the initial weight of the reactor. The tests were terminated after 331 days.

#### 2.3.2 Benthic test (seafloor scenario)

The biodegradation under aerobic conditions at the water-sediment interface was tested following Tosin et al. (2012) and ISO 19679 (2016). At the start of the experiments, the medium used for the benthic lab test had a pH of 8.3, a water content of 95.8%, a total solids content of 4.2%, of which volatile solids were 15.7%. The tests were carried out in 250 ml Erlenmeyer flasks with a container attached to a side port filled with 3 mL 0.5 N KOH for CO_2_ absorption. The bottle and the container were closed with a Python rubber stopper (Rubber BV, Hilversum, The Netherlands). The bottles were incubated in the dark in a closed box that was placed in a temperature-controlled room (20 ± 1 °C). The test materials (PBSe and PBSeT) were square-shaped specimens with a dimension of approximately 30 × 30 mm. The negative control was done with a specimen of LDPE. For the test with PHA as the positive control specimens were cut in circles of 20 mm diameter. The mass of each specimen (about 36 mg for PHA, about 25 mg for the others) was recorded. For the test 15 reactors were prepared to enable testing in triplicate of PBSe, PBSeT, PHA (reference material/positive control, LDPE (negative control), and a blank without test material to correct for endogenous respiration. Sediment (30 g) was placed at the bottom of each reactor with 70 ml seawater and 3 ml of 0.5N KOH (the CO_2_ absorbing solution) was introduced in the container. Endogenous respiration was measured after one week by measuring the CO_2_ production. The results showed that the endogenous respiration was similar in the bottles. After the initial week of pre-incubation, the test was started. For this, the reactors were opened and the specimen of test materials PBSe or PBSeT or LDPE or PHA were placed on top of the sediment. About 25-35 mg of test material or LDPE or PHA was introduced in the reactors. The blanks for endogenous respiration did not receive any test specimen. The carbon dioxide produced in each reactor reacted with KOH and was titrated as described above. The tests were run for 331 days. The content of the test bottles was sacrificed for the retrieval of remaining plastic test items after the termination of the test.

#### 2.3.3 Modelling the half-life *t*_0.5_ of biodegradation

For all statistical analyses, R project was used (R Core Team, 2020). From the raw data of CO_2_ release over time (equaling aerobic biodegradation) the half-life of the polymer was analyzed using Three Parameter Logistic Regression (3PL) by fitting the data with a non-linear model (package *nlme*) to the formula:

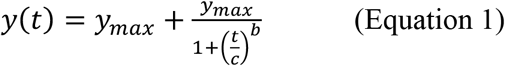

where *y* is the polymer biodegradation in percent, *t* the corresponding time in days, and *y*_max_, *b,* and *c* the curve parameters estimated by the model and representing the biodegradation at the plateau, the slope-factor and the inflection-point of the curve, respectively (adapted from Junker et al. 2016). Since the same test flask was measured repeatedly over time, a random factor was included to estimate *y*_max_. If no model parameters could be estimated, *y*_max_ was below 0 (as for LDPE), and/or estimated parameters (except *y*_0_) were not significantly different to 0 no biodegradation was assumed. To visualize the confidence interval (CI) of the predicted disintegration curves, a Monte Carlo simulation approach was employed to re-sample for each *x* value (days) 50,000 predicted values (% biodegradation) considering the estimates and variance of model parameters. The range of the 95 % percentile of these samples was considered as 95 % confidence interval. Before calculating the half-life, it had to be tested if *y*_max_ was higher than 50 %. Therefore, empirical p-values were used. 500,000 values for *y*_max_ were re-sampled considering the coefficients predicted by the non-linear model and its variance. If less than 5 % of these values were below 50 %, the plateau was assumed to be greater than or equal to 50 %. If this condition was fulfilled, 500,000 values for the half-life (*t*_0.5_) were generated by setting *y* in the 3PL equation to 50 %, using the estimated values and corresponding variances of the model parameters, and solving the formula to *t*. In some cases, the half-life could not be calculated as described because *ymax* assumed values were below 50 % (however in less than 5 % of the cases, as shown by the previous test). In these cases, the half-life was set to infinity. To compare calculated values for *t*_0.5_ between habitats and experiments, the difference between two groups of *t*_0.5_ values was calculated. For these pairwise comparisons, empirical *p*-values were calculated 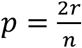 where *r* is the number of *t*_0.5_ differences either above or below zero and *n* the number of trails (i.e. 500,000). If all *r* values were below or above 0, the lowest possible value for *p* was assumed (2·1/*n* = 4·10^−6^). *p*-values were adjusted for multiple comparisons using the Holm method (Holm, 1979). The distributions of the re-sampled half-life values were visualized by violin plots in which the spread along the *x* axis represents the frequency of values on the *y* axis (half-life). Different letters indicate significantly different groups.

### 2.4 Mesocosm tests

The mesocosm tests were conducted in two consecutive years (y1 and y2) in triplicates in a climate chamber as described before (Lott et al., 2020). Three coastal habitats were simulated in a tank system consisting of two 630-L plastic containers placed on top of each other. The upper tank contained a layer of siliclastic beach sediment in which the samples were buried (eulittoral scenario) and which was flooded with seawater every 12 h. The bottom of the upper tank was perforated to allow the water to slowly drain to the lower tank, thus mimicking a 12-h tidal cycle at the samples. The lower tank contained a layer of carbonate seafloor sediment on which samples for the benthic test were placed. The pelagic test was performed by hanging samples in the water column of the lower tank, which was illuminated on a 12:12 h rhythm. The bulk water of the two tanks was connected by pipes and constantly moved by additional pumps in the lower tank. The water was regularly checked for salinity, pH, and oxygen saturation and compensated for evaporation loss by adding demineralized water if necessary. The temperature was 20.5 °C ± 1 °C and mean light intensity on the sediment surface of the benthic tests was 11.56 μmol photons·m^−2^·s^−1^. Salinity was 39 ± 1. The pH was 8.1 ± 0.1. The oxygen concentration was close to air saturation (98 ± 2 %). In the first year (y1), three polymers were tested in the mesocosm experiments: LDPE, PHA, and PBSeT were sampled at four time points (t1 – t4) (SI Table 1). In the second year (y2), PBSe was tested additionally and sampled at two time points (t1 and t3). Three to five samples were retrieved ca. every 2.5 months from the tanks. The last sampling interval of the y2 experiment was only 1.5 months.

### 2.5 Field tests

Field tests were performed in the eulittoral (beach scenario), the pelagic (open water scenario), and the benthic (sublittoral seafloor) (Figure 1) as described before (Lott et al., 2020). For environmental conditions at each site see Table 2. For sampling dates and intervals see SI Table 2.

#### 2.5.1 Eulittoral tests

The eulittoral tests were set up on the Island of Elba, Terme di San Giovanni, Portoferraio (N 42°48’12.1“N, 010°19’01.0“E, Italy in a former saline basin, now open to the sea (Lott et al., 2020). The test system consisted of 60-L plastic bins filled with beach sediment in which the samples were buried. To simulate an intertidal sandy beach the bins were placed on wooden racks in the midwater line in a way that the samples were exposed to changing conditions of being wetted and falling dry with the tides.

#### 2.5.2 Pelagic and benthic tests

The pelagic and benthic field tests in the Mediterranean Sea were performed in the marine protected area of the National Park Tuscan Archipelago off the Island of Pianosa (42°34’41.4“N, 010°06’30.6“E), Italy. The benthic field tests in South-East Asia were performed in Sahaong Bay (01°44’35.4”N, 125°09’09.3”E), Pulau Bangka, NE Sulawesi, Indonesia. For details see Lott et al. (2020).

The pelagic test systems consisted of a rack to which the sample frames were attached, anchored to stay afloat at a water depth of 20 m, chosen to avoid influence by surface wave activity. For the benthic tests, the samples were mounted to a flat panel which was fixed to the seafloor at 40 m in the Mediterranean Sea and to 32 m in Indonesia. These depths were chosen to avoid interference of the experiments with the seagrass meadows or coral reef structures respectively. In the Mediterranean field tests, in the first run of field experiments (y1) in all three habitats five replicates of PHA, PBSeT, and LDPE were exposed and sampled about every 2.5 months. One additional set was left exposed for two years (t_5_), together with new samples for a second 1-year run (SI Table 2). From these additional sets, two replicates were sampled from the pelagic and benthic experiment after 678 days and five replicates were sampled from the eulittoral test system after 686 days. In the second experiment run (y2), the number of replicates per time was reduced to three. Also, PBSe was added as test material with three replicates in the pelagic and benthic tests, and two replicates per time in the eulittoral test. The data from y1 and y2 were pooled in the analysis as experiment y1 followed the same seasonality as y2.

In the wet tropics of South-East Asia, tests were conducted in 2017 and 2018. Three replicates of PHA, PBSe, and PBSeT were exposed to the benthic habitat. Sampling occurred after 19, 48, 62, 90, 310 days for PHA, after 16, 90, 130, 310 days for PBSe and after 90 and 310 days for PBSeT.

### 2.6 Sampling and data acquisition from mesocosm and field tests

At the given time interval, the sample frames were carefully detached from their racks (pelagic, benthic) or dug out of the sediment (eulittoral), rinsed in ambient water, packed singly in plastic (PE) bags covered with water, and brought to the laboratory for further treatment the same day. Each frame was opened and the sample photographed in a standardized way. Then the samples were washed in freshwater and left to dry at room temperature overnight.

#### 2.6.1 Disintegration measurements

In the open-system tests of the mesocosm and field experiments, material disintegration was determined as a proxy for biodegradation assuming that the samples were protected from mere physical damage well enough by the construction of the test frame. The degree of disintegration (% area loss) of each sample was determined photogrammetrically. Dried samples (Mediterranean tests) were scanned on a LIDE 210 flatbed scanner (Canon Inc.). For the Indonesian samples, photos (Canon EOS 5D MkII) of the freshly sampled films were used and analyzed for the proportion of lost vs. still intact surface using the software ImageJ (https://imagej.nih.gov/ij/).

#### 2.6.2 Calculating half-life *t*_0.5_ based on disintegration data

Since in the field and mesocosm experiments far fewer data points than in the lab experiment were acquired, a model with fewer degrees of freedom (fewer estimated parameters) had to be chosen. The disintegration of the polymer area over time was analyzed using beta regression (package *betareg*) because response values consisted of percent data. The extreme values of 0 % and 100 %, were transformed according to Smithson and Verkuilen (2006). For the beginning of the experiment, no disintegration was assumed, therefore artificial values with 0 % disintegrated area were included. For each habitat and experiment (field vs. mesocosm) one model was applied. For the mesocosm experiment, additionally, the influence of the year of the experiment was analyzed and integrated into further analysis if a significant interaction with the variable time existed. Different models were applied using logit, cloglog, cauchit, and loglog link-functions (see SI Table 3). The best model was then selected by comparing the Root Mean Square Deviation (package *caret*). When comparing the disintegration rate between habitats it is not sufficient to only compare the slopes. The non-linear shape of the back-transformed disintegration curve depends on both, the *y*-intercept *a* and the slope *b*. Therefore, rather half-life (*t*_0.5_) should be compared. Since different link-functions were used, different formulas had to be applied to calculate the corresponding *x* (time) values at which the disintegration reached 50 %. For example, in the case of *logit*-transformed data, 50 % of the material is left at the *x*-intercept of the regression curve for transformed data:

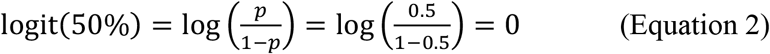

**Table 3:**
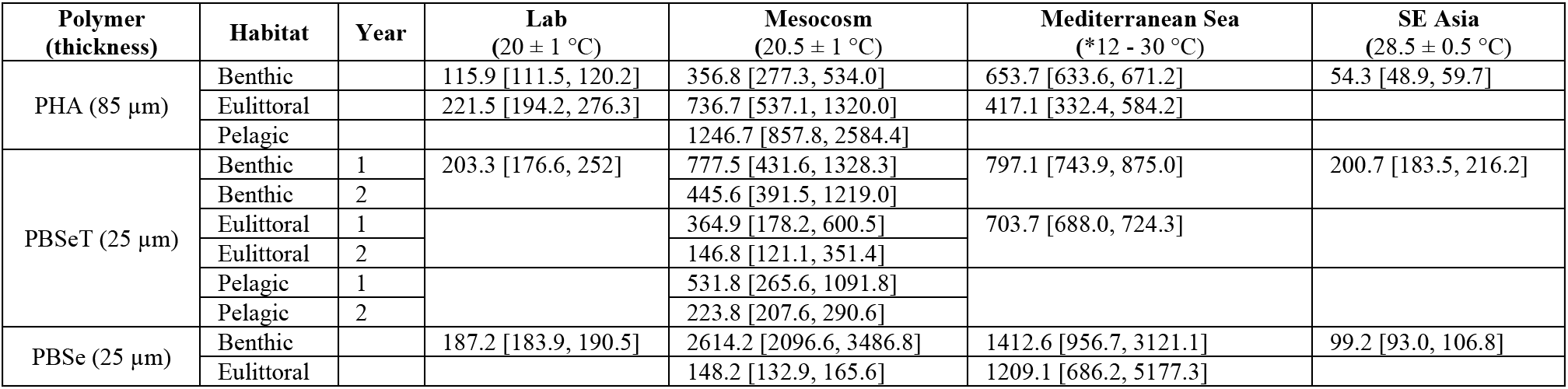
Summary of all tests: Half-lives *t*_0.5_ of PHA, PBSe, and PBSeT as predicted by statistical modelling of the experimental data. The range of the confidence interval CI (centered 95-percentile of the Monte Carlo simulations) is given in brackets. * = temperature range (daily mean): pelagic 14 - 25 °C, benthic 14 - 20 °C, eulittoral 12 - 30 °C

| Polymer (thickness) | Habitat | Year | Lab (20 ± 1 °C) | Mesocosm (20.5 ± 1 °C) | Mediterranean Sea (*12 - 30 °C) | SE Asia (28.5 ± 0.5 °C) |
| --- | --- | --- | --- | --- | --- | --- |
| PHA (85 $\mu\text{m}$) | Benthic | | 115.9 [111.5, 120.2] | 356.8 [277.3, 534.0] | 653.7 [633.6, 671.2] | 54.3 [48.9, 59.7] |
|  | Eulittoral |  | 221.5 [194.2, 276.3] | 736.7 [537.1, 1320.0] | 417.1 [332.4, 584.2] |  |
|  | Pelagic |  |  | 1246.7 [857.8, 2584.4] |  |  |
| PBSeT (25 $\mu\text{m}$) | Benthic | 1 | 203.3 [176.6, 252] | 777.5 [431.6, 1328.3] | 797.1 [743.9, 875.0] | 200.7 [183.5, 216.2] |
|  | Benthic | 2 |  | 445.6 [391.5, 1219.0] |  |  |
|  | Eulittoral | 1 |  | 364.9 [178.2, 600.5] | 703.7 [688.0, 724.3] |  |
|  | Eulittoral | 2 |  | 146.8 [121.1, 351.4] |  |  |
|  | Pelagic | 1 |  | 531.8 [265.6, 1091.8] |  |  |
|  | Pelagic | 2 |  | 223.8 [207.6, 290.6] |  |  |
| PBSe (25 $\mu\text{m}$) | Benthic | | 187.2 [183.9, 190.5] | 2614.2 [2096.6, 3486.8] | 1412.6 [956.7, 3121.1] | 99.2 [93.0, 106.8] |
|  | Eulittoral |  |  | 148.2 [132.9, 165.6] | 1209.1 [686.2, 5177.3] |  |

The *x*-intercept can be calculated by 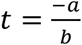. The formulas used for the other link-functions are compiled in SI Table 3. A Monte Carlo simulation approach was applied to re-sample 500,000 values for *a* and *b* considering model parameter estimates and variance and a normal distribution. The comparison of the generated half-life was performed with empirical *p*-values as described for the lab experiment.

## 3 Results

### 3.1 Laboratory tests with Mediterranean Sea matrices

The lab tests with matrices from two of the shallow water habitats chosen for field and mesocosm tests were done to prove the biodegradability of the three tested polymers PHA, PBSe, and PBSeT under optimized lab conditions with natural matrices. The eulittoral test (beach scenario) and the benthic test (seafloor scenario) were selected because the tested polymers have a higher specific density than seawater and will sink to the seafloor. The pelagic test was not performed due to space limitations in the laboratory. The biodegradation in both habitats was fastest for PHA and similar for PBSeT and PBSe (Figure 2).

In the benthic test, a plateau was reached or nearly reached after about one year for the three test materials. PHA was more or less completely converted to CO_2_ as is evidenced by the 81% biodegradation that was calculated from the CO_2_ production. The biodegradation percentage of PBSe and PBST was 71% and 76%, respectively, which also indicates that mineralization of these test items was almost complete, given that an incorporation of about 25 – 30 % of substrate carbon into microbial biomass is assumed (see e.g. Payne (1970) for conversion rates in bacterial cultures). Except for LDPE none of the plastic test items could be retrieved, not even in fragments, from the benthic test bottles which indicates that the disintegration of the test items was complete. In some cases, e.g. for PBSe very small particles were visible but these were in the size range of sand grains. LDPE was recovered intact from the flasks and biodegradation was not detected. In the eulittoral test, none of the three materials reached the plateau phase within the duration of the experiment (331 d). PHA was completely disintegrated in all flasks. For both, PBSe and PBSeT, one of the three replicates could be retrieved degraded to 25 and 33%, respectively.

The respective half-lives *t*_0.5_ predicted by the model are given in the following text without the confidence intervals which can be found as a summary in Table 3 at the end of the results section. For PHA in the benthic test, the half-life was 116 d and significantly (*p* < 0.001) lower than in the eulittoral test (222 d, Figure 3). For PBSe, the modelled *t*_0.5_ was 203 d and for PBSeT 187 d both in the benthic test. In the eulittoral test, the exposure time for both materials was not sufficient to reach a biodegradation of more than 50 %, therefore modelling *t*_0.5_ was not possible.

**Figure 3.**
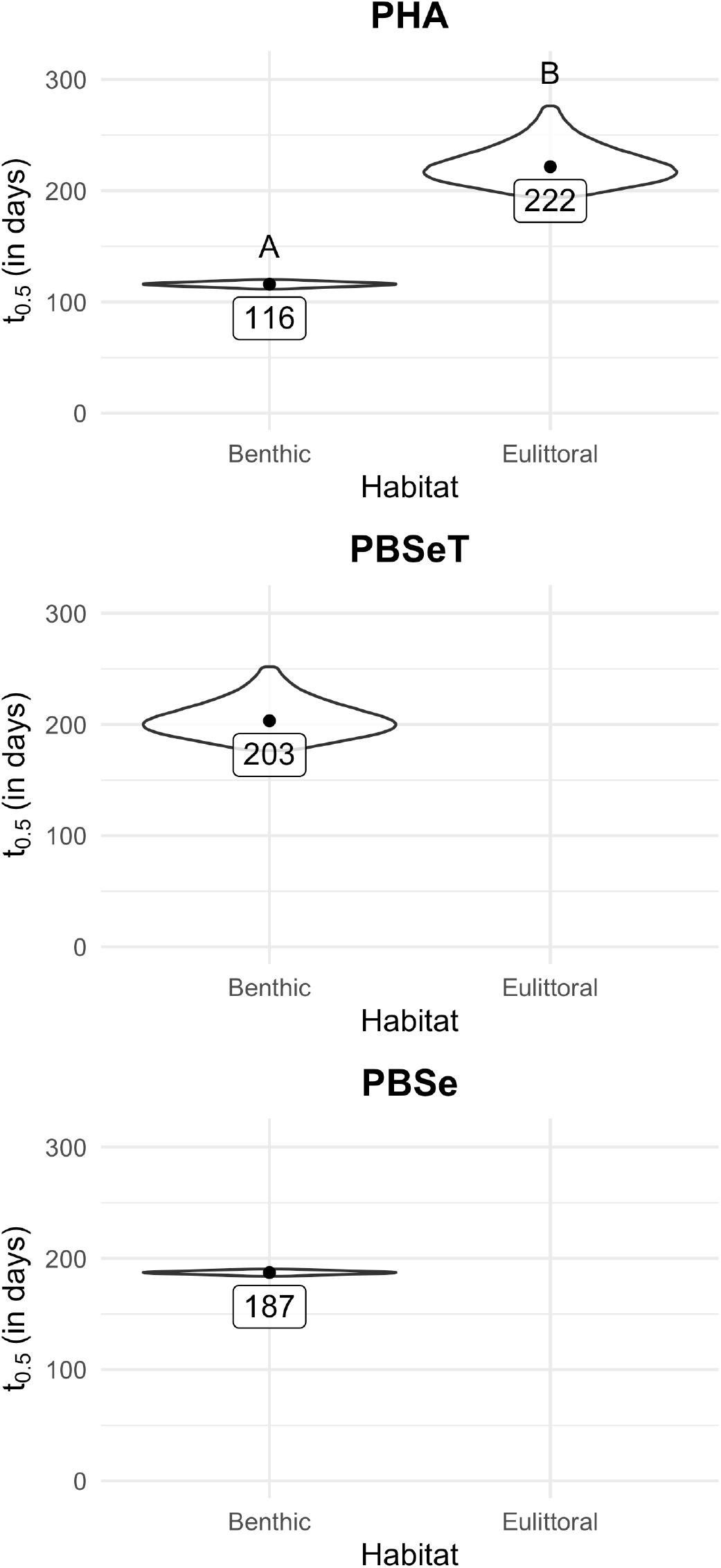
Laboratory experiments with Mediterranean Sea water and sediment: Biodegradation half-lives *t*_0.5_ of PHA, PBSe, and PBSeT. Monte Carlo simulations of half-life for tested polymers in two habitats, on the seafloor at the sediment-seawater interface (benthic) and the beach scenario (eulittoral). Shown is the distribution of the centered 95 percentiles of the simulations as violin plots. The dot in the center of each violin represents the half-life as calculated from the estimated model coefficients; the value is given in the box below. Letters above the violin plot represent the results of the multiple comparisons. Different letters (A, B) indicate significantly different groups (i.e. Holm-adjusted *p*-value below *α* = 0.05). For PBSeT and PBSe in the eulittoral habitat, no half-life could be calculated because the plateau phase was not reached and values remained below 50% (*cf.* fig. 2).

### 3.2 Mesocosm tests with Mediterranean Sea matrices

All test materials, except LDPE, showed disintegration with high heterogeneity between replicates, habitats, and material type. About 90 % disintegration was observed for PBSe and PBSeT in the eulittoral test in y2 after 238 days (Figure 4) and for PBSeT in the pelagic test in y2 after 271 days. Most samples of PBSe and PBSeT in the other habitats were disintegrated less than 50 % after 308 and 270 days of exposure, respectively.

**Figure 4.**
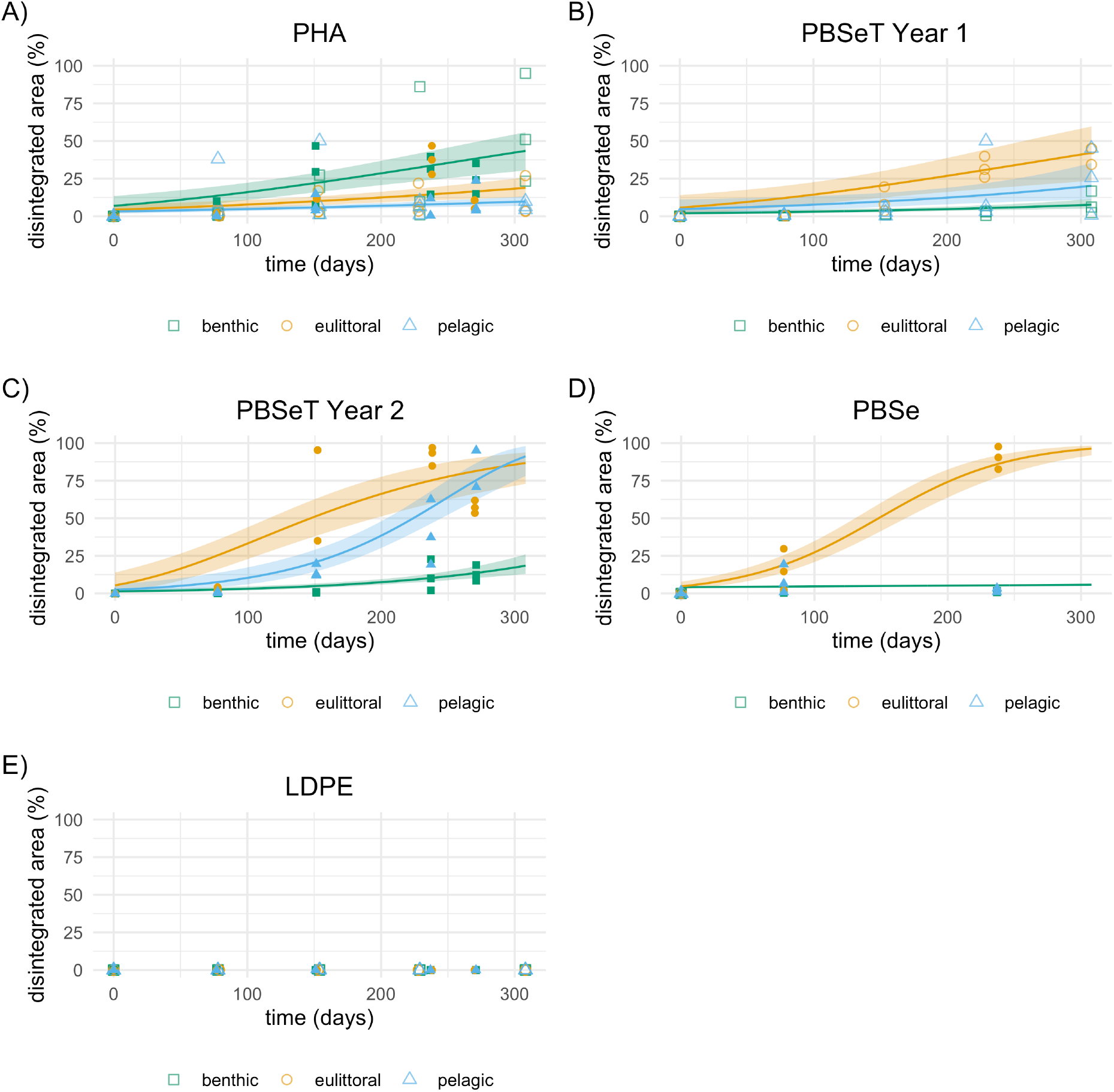
Mesocosm experiments with Mediterranean Sea water and sediment: Disintegration curves of PHA, PBSeT, and PBSe. Disintegrated area of polyhydroxyalkanoate (PHA) as positive control, test materials polybutylene sebacate co-terephthalate (PBSeT) and polybutylene sebacate (PBSe) and low-density polyethylene (LDPE) as negative control over 10 (year 1) and 9 (year 2) months when exposed in the seafloor scenario at the sediment-seawater interface (benthic), in the water column (pelagic) scenario or buried in the intertidal sandy beach (eulittoral) scenario. The model curve ± SE (shaded area) was drawn if the slope was different from zero and positive. Dots represent raw data analyzed in the model (open = year 1, filled = year 2), For PBSeT two significantly different model curves are shown. Colors represent the habitat (green = benthic, orange = eulittoral, blue = pelagic).

The half-life of PHA did not differ between the two experiments (year 1 and year 2) within each habitat. The predicted half-life in the benthic habitat (357 d) was significantly lower than in the eulittoral (737 d) and pelagic (1247 d) habitats (Figure 5, Table 3, SI Table 4). PBSeT had a *t*_0.5_ of 778 d in y1 and 446 d in y2 in the benthic tests, 365 d in y1 and 147 d in y2 in the eulittoral tests, and 532 d in y1 and 224 d in y2 in the pelagic tests (Figure 5, Table 3). In the eulittoral experiment, the degradation was significantly faster in the second-year experiment than in the first-year experiment (*p* = 0.0071). No significant differences were detected between both experiments (y1 and y2) in the benthic and pelagic tests (SI Table 5). The half-life of PBSe was significantly higher in the benthic tests (2614 d) than in the eulittoral test (148 d, *p* < 0.0001, Figure 5, Table 3). Predictions for the half-life of PBSe in the benthic test should be considered with caution, since maximum degradation after 237 d was 1.5 %, making predictions of the time needed until 50 % is degraded unreliable. In the pelagic test, no significant disintegration was measured for PBSe. For LDPE no disintegration at all was measured in any test during the exposure time thus it was not possible to model *t*_0.5_.

**Table 4:**
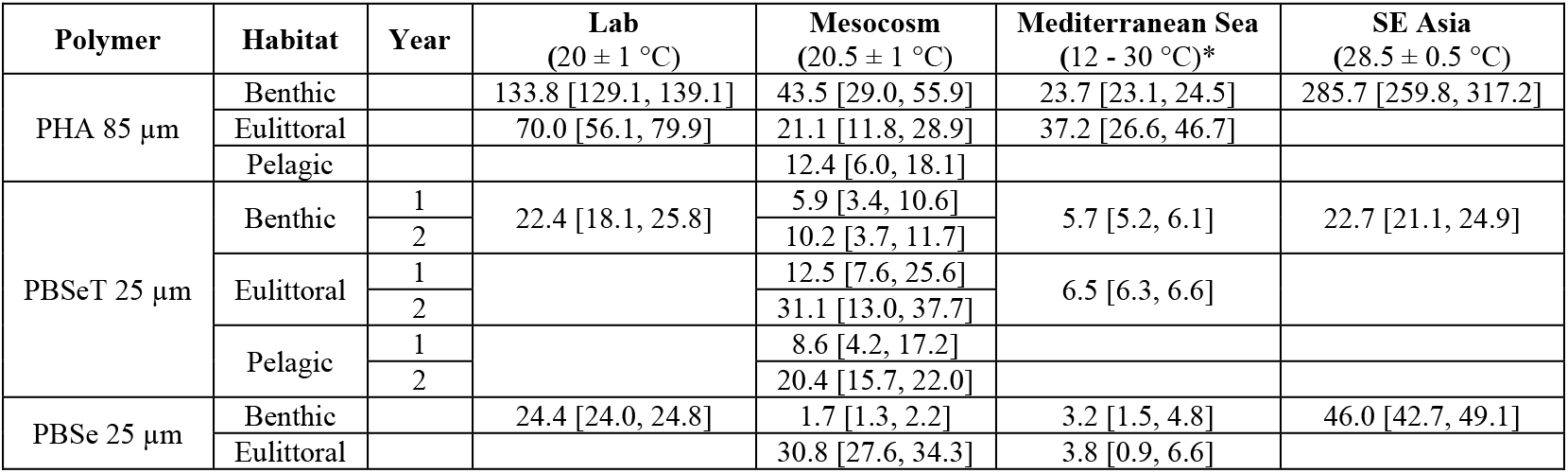
Surface micro-bioerosion rates (μm · yr^−1^) of PHA, PBSe, and PBSeT under different marine conditions. The range of the confidence interval CI (centered 95-percentile) is given in brackets. * = temperature range (daily mean): benthic 14 - 20 °C, eulittoral 12 - 30 °C.

| Polymer | Habitat | Year | Lab<br>($20 \pm 1 \text{ }^\circ\text{C}$) | Mesocosm<br>($20.5 \pm 1 \text{ }^\circ\text{C}$) | Mediterranean Sea<br>($12 - 30 \text{ }^\circ\text{C}$)* | SE Asia<br>($28.5 \pm 0.5 \text{ }^\circ\text{C}$) |
| --- | --- | --- | --- | --- | --- | --- |
| PHA 85 $\mu\text{m}$ | Benthic | | 133.8 [129.1, 139.1] | 43.5 [29.0, 55.9] | 23.7 [23.1, 24.5] | 285.7 [259.8, 317.2] |
|  | Eulittoral |  | 70.0 [56.1, 79.9] | 21.1 [11.8, 28.9] | 37.2 [26.6, 46.7] |  |
|  | Pelagic |  |  | 12.4 [6.0, 18.1] |  |  |
| PBSeT 25 $\mu\text{m}$ | Benthic | 1 | 22.4 [18.1, 25.8] | 5.9 [3.4, 10.6] | 5.7 [5.2, 6.1] | 22.7 [21.1, 24.9] |
|  |  | 2 |  | 10.2 [3.7, 11.7] |  |  |
|  | Eulittoral | 1 |  | 12.5 [7.6, 25.6] | 6.5 [6.3, 6.6] |  |
|  |  | 2 |  | 31.1 [13.0, 37.7] |  |  |
|  | Pelagic | 1 |  | 8.6 [4.2, 17.2] |  |  |
|  |  | 2 |  | 20.4 [15.7, 22.0] |  |  |
| PBSe 25 $\mu\text{m}$ | Benthic | | 24.4 [24.0, 24.8] | 1.7 [1.3, 2.2] | 3.2 [1.5, 4.8] | 46.0 [42.7, 49.1] |
|  | Eulittoral |  |  | 30.8 [27.6, 34.3] | 3.8 [0.9, 6.6] |  |

**Figure 5.**
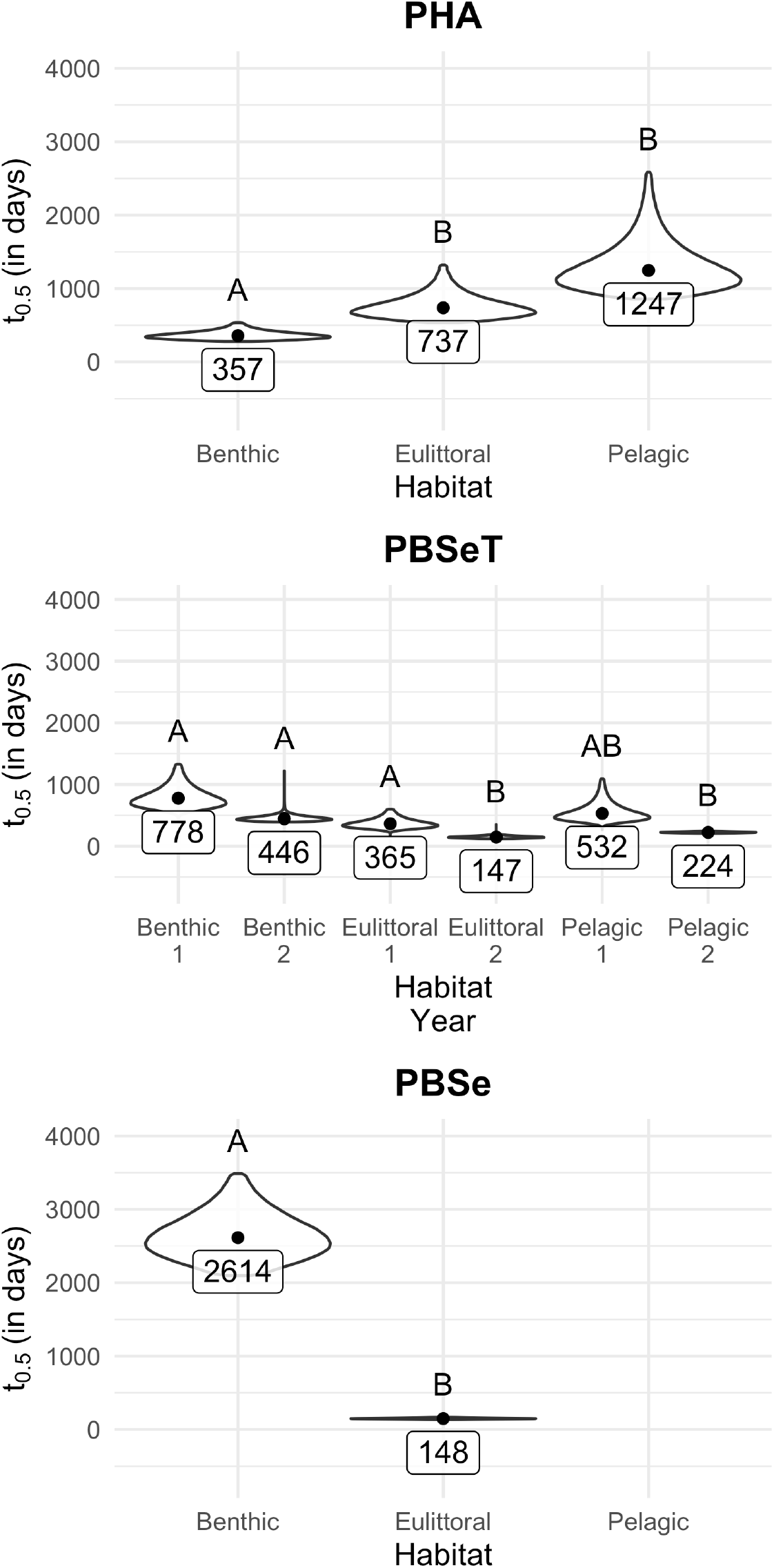
Mesocosm experiments with Mediterranean Sea water and sediment: Disintegration half-lives *t*_0.5_ of PHA, PBSeT, and PBS. Monte Carlo simulations of half-life for tested polymers in three habitats (on the seafloor at the sediment-seawater interface (benthic), the beach (eulittoral), and the water column (pelagic) scenario). Shown is the distribution of the centered 95 percentiles of the simulations as violin plots. The dot in the center of each violin plot represents the half-life as calculated from the model coefficients, and the value is given in the box below. Letters above the violin plot represent the results of the multiple comparisons. Different letters indicate significantly different groups.

### 3.3 Field tests

#### 3.3.1 Mediterranean Sea

All test materials, except LDPE, showed signs of disintegration when in contact with sediment, i.e. in eulittoral and benthic tests (Figure 6). In the pelagic tests, no disintegration was observed for PHA and PBSe within 2 years. For PBSeT, material disintegration increased significantly over time, but at a very low slope. The maximum disintegration after 676 d was only 1.01 % and 0.13 %, therefore we did not predict the half-life. The heterogeneity between replicates was high. The half-life *t*_0.5_ of PHA in the benthic test (654 d) was significantly higher (*p* = 0.0161) than in eulittoral test (417 d, Figure 8, Table 3, SI Table 6). In the pelagic test, modeling of *t*_0.5_ was not possible because no disintegration was measured during the exposure time. PBSeT had a *t*_0.5_ of 797 d in the benthic test, which was significantly lower than in the eulittoral test (704 d, Figure 8, Table 3, SI Table 7). PBSe had a *t*_0.5_ of 1413 d in the benthic test and of 1209 d in the eulittoral test (Figure 8, Table 3, SI Table 8).

**Figure 6.**
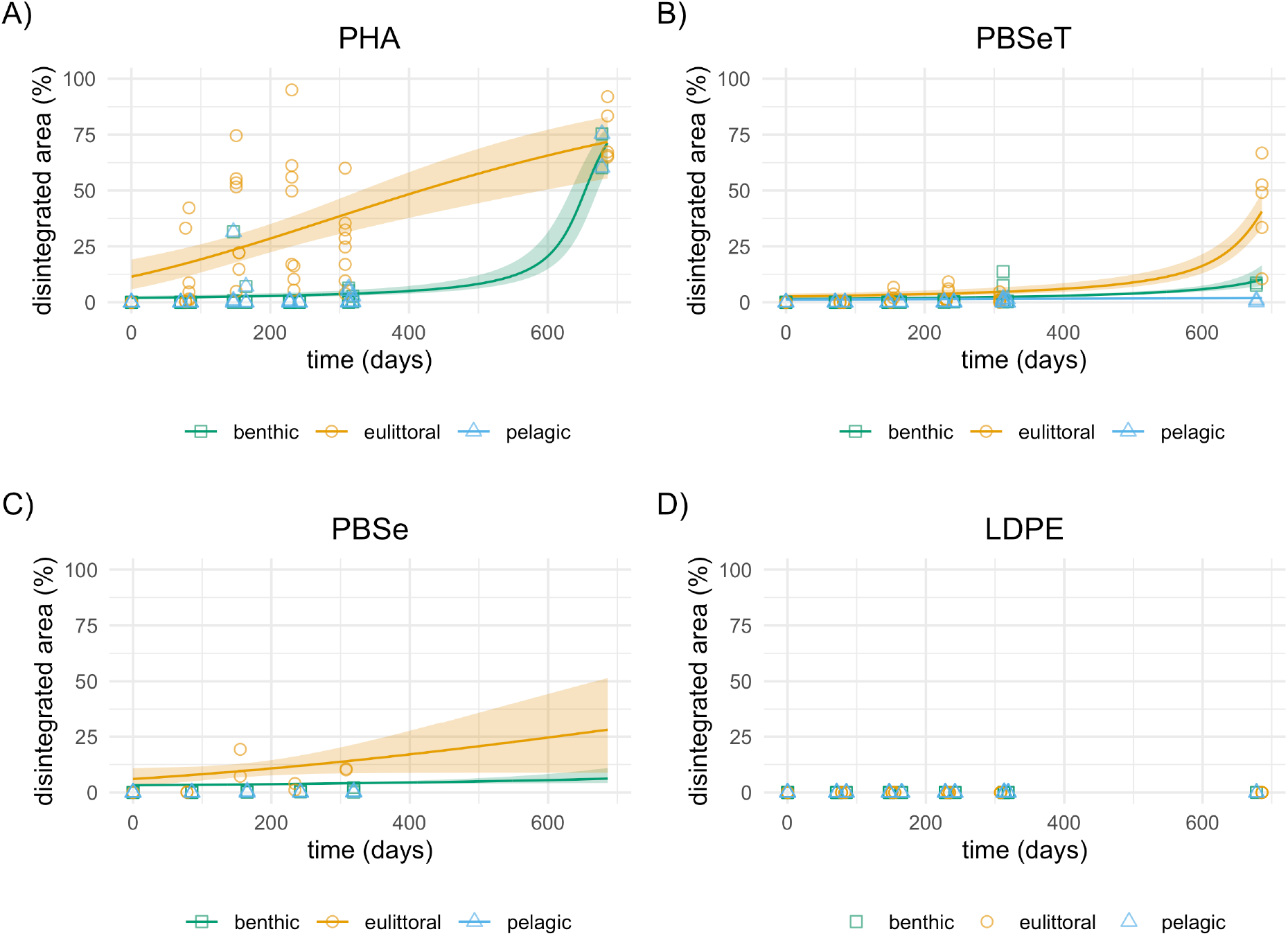
Field experiments in the Mediterranean Sea: disintegration curves of PHA, PBSeT, and PBSe: Disintegrated area of polyhydroxyalkanoate (PHA) as positive control (A), test materials polybutylene sebacate co-terephthalate (PBSeT, B) and polybutylene sebacate (PBSe, C) and low-density polyethylene (LDPE, E) as negative control over 10 (y1) and 9 (y2) months and PBSe over 9 months (y2) when exposed on the seafloor at the sediment-seawater interface (benthic), the beach (eulittoral) and the water column (pelagic) scenario. The model curve ± SE (shaded area) was drawn if the slope was different to zero and positive. Dots represent raw data analyzed in the model. Colors represent the habitat (green = benthic, orange = eulittoral, blue = pelagic).

#### 3.3.2 SE Asia

All test materials, except LDPE, fully disintegrated in the benthic test within few months (Figure 7) with heterogeneity between replicates. The *t*_0.5_ for PHA was 54 d, for PBSeT 201 d, and for PBSe 99 d (Figure 8, Table 3). The disintegration of all polymer types was significantly faster in SE Asia compared to the Mediterranean Sea (Figure 8, SI Tables 6 - 8). Summarized (Table 3), in the field tests in the Mediterranean Sea, the *t*_0.5_ of a 25 μm thick film of PBSeT and PBSe and an 85 μm thick film of PHA ranged between 1.8 - 3.9 years in the benthic tests (seafloor scenario) and 1.1 - 3.3 years in the eulittoral tests (beach scenario). No significant disintegration occurred for any of the materials in the pelagic tests (water column scenario). In the benthic test in SE Asia, *t*_0.5_ was about 0.15 years (2 months) for PHA, 0.55 years (7 months) for PBSeT, and 0.27 years (4 months) for PBSe. For LDPE no disintegration was measured, thus no half-life was possible to calculate.

**Figure 7.**
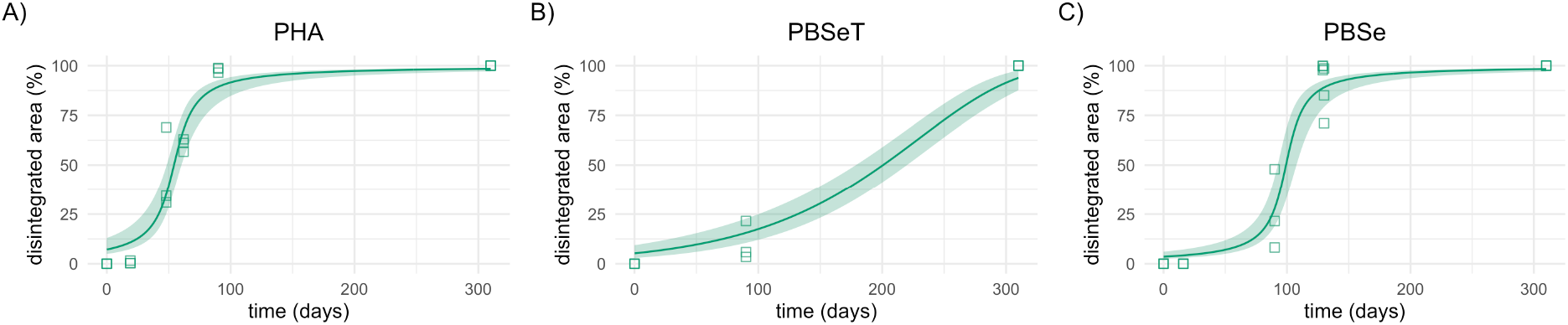
Field experiments in tropical SE-Asia: Disintegration curves of PHA, PBSeT, and PBSe. Disintegrated area of tested polymers (A) PHA, B) PBSeT, C) PBSe) for almost one year (310d) when exposed on the seafloor sediment-seawater interface (benthic). The model curve ± 95% CI (shaded area) was drawn if the slope was different from zero and positive. Points represent raw data analyzed.

**Figure 8.**
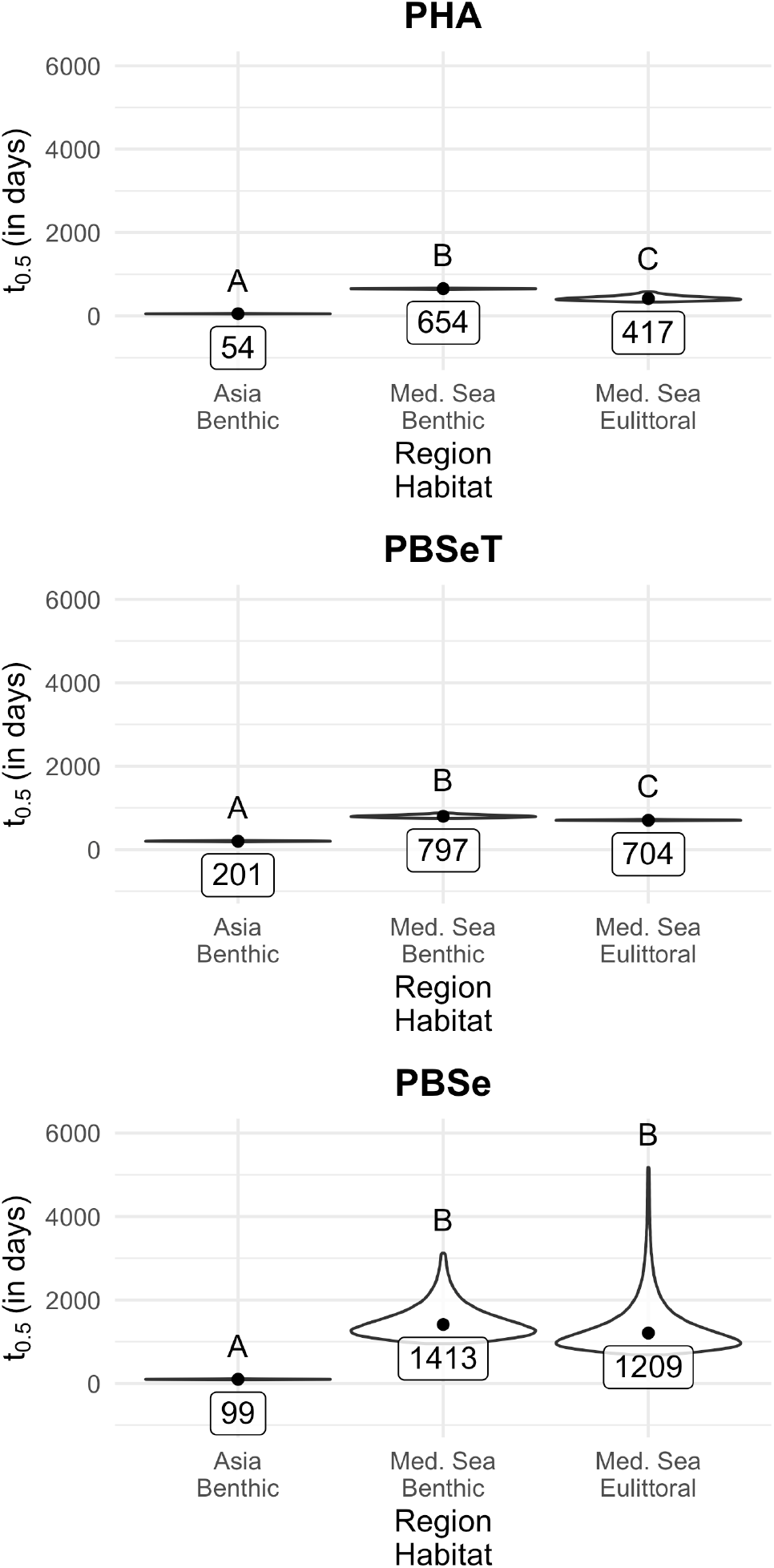
Comparison of the disintegration half-lives *t*_0.5_ of PHA, PBSeT, and PBSe from field experiments in tropical SE Asia and the Mediterranean Sea: Monte Carlo simulations of half-life for tested polymers in the benthic (seafloor sediment-seawater interface) in Asia and the Mediterranean Sea and in the eulittoral (beach scenario) in the Mediterranean Sea. Shown is the distribution of the centered 95 percentiles of the simulations as violin plots. The dot in the center of each violin plot represents the half-life as calculated from the model coefficient and its value is given in the box below. Letters above the violin plot represent the results of the multiple comparisons. Different letters indicate significantly different groups (i.e. Holm-adjusted *p*-value below *α* = 0.05).

**Figure 9.**
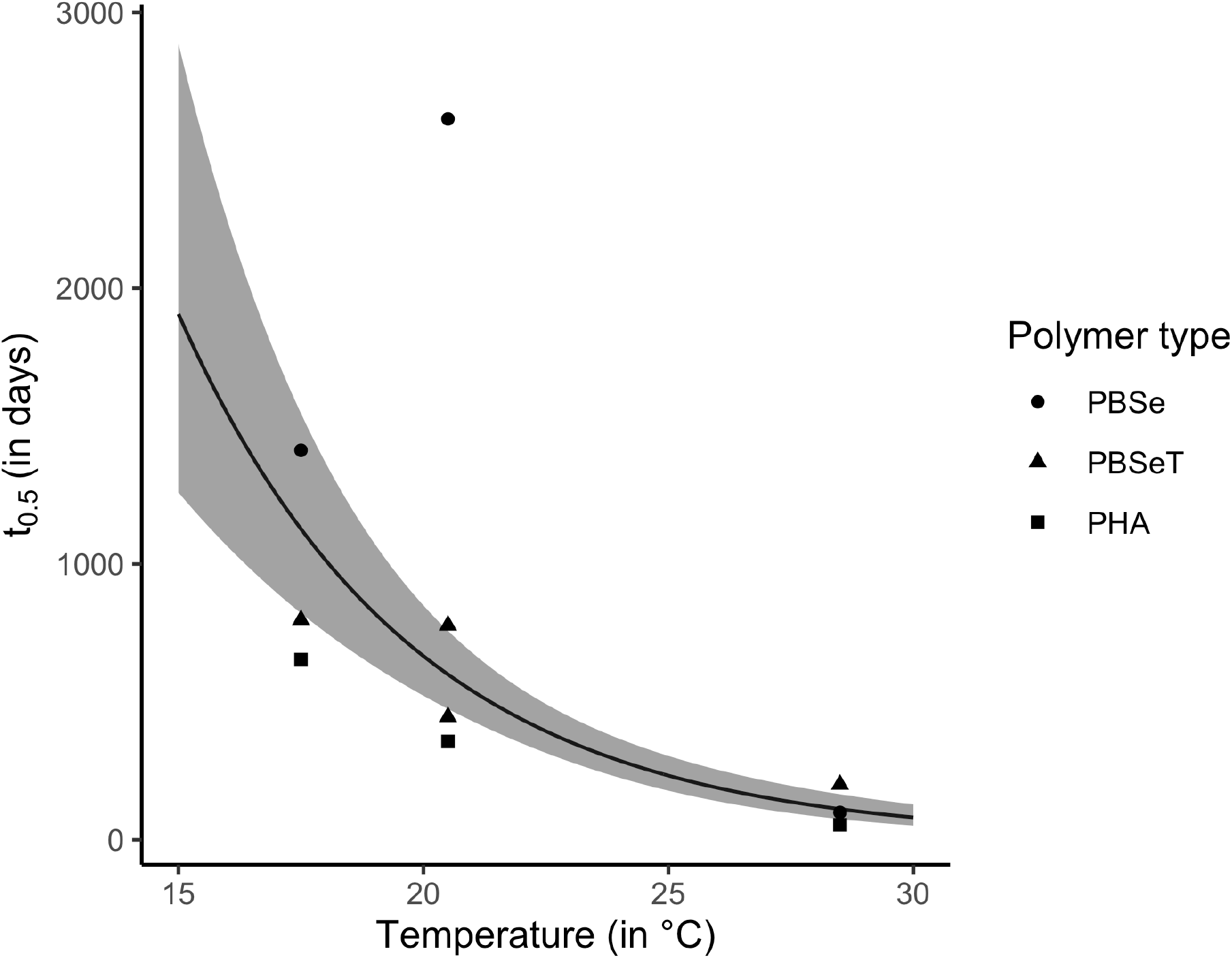
Temperature dependence of half-life in benthic tests: half-life *t*_0.5_ decreases exponentially with temperature. The dots are the predicted half-life values for PHA (squares), PBSeT (triangles), and PBSe (circles). The line represents an exponential model, the shaded area is the standard error.

## 4 Discussion

The aim of this study was to prove the biodegradability of biodegradable polymers under laboratory conditions, to test the biodegradation performance under natural field conditions, and in a tank test system with natural matrices.

We tested the performance of PHA (as reference material and positive control), PBSe, and PBSeT in three different habitat scenarios, namely the intertidal sandy beach (eulittoral), the sandy sublittoral seafloor (benthic), and the open water column (pelagic) in a three-tier test approach in closed-vessel laboratory, in mesocosm tank and in-situ field tests, some of which in two different climate zones (the warm-temperate Mediterranean Sea and the tropical sea of SE Asia).

Furthermore, we introduced an analytical tool based on statistical modelling to predict the specific half-life calculated from the experimental data for each set of test conditions, to use the half-lives to numerically compare the performance of three different materials tested, and to numerically compare the performance of one material in different environmental settings. The three-tier approach to test the performance of biodegradable plastic in the marine environment gave a comprehensive view and could differentiate between sediment type, habitat, and climate zone.

### 4.1 Half-lives *t*_0.5_ differed between the materials in all test systems and habitats

#### 4.1.1 Laboratory experiments

In the lab tests, biodegradation of PHA was faster in the benthic than in the eulittoral test. For PBSe and PBSeT in the eulittoral tests the plateau phase of biodegradation, as calculated by CO_2_ evolution, was not reached within the test period and values remained below 50 %. Consequently, *t*_0.5_ could not be calculated. The levelling out of the biodegradation curves in the benthic test (Figure 2) indicates a limitation of substrate for the bacteria and the nearly complete conversion of the polymer into CO_2_ and microbial biomass. In the humid sand of the eulittoral tests, the CO_2_ production rate was generally lower than at the submersed sand surface in the benthic test. However, the fact that in most of the experiments no or only little test material was left could indicate a more efficient build-up of microbial biomass in the eulittoral, a (yet) incomplete mineralization of soluble intermediates of biodegradation, a limitation of nutrients or inhibition by a hypothetical starvation factor as proposed by Mistriotis et al. (2019) in soil. For PHA, both lab tests applied showed the biodegradability of the positive control but also revealed significant differences in the performance under the different experimental conditions of a simulated seafloor (benthic) and a simulated beach (eulittoral). The lab tests were conducted under static conditions without stirring or shaking the medium leaving the system purely diffusion-driven. It can be assumed that oxygen availability to the acting microbes was most of the time limited. This is also corroborated by the observation of black spots on the sediment below some of the samples in the benthic test indicating the precipitation of dark metal salts under anoxic conditions. As the methods for both tests are defined as aerobic (Tosin et al., 2012; ISO 22404, 2019; ISO 19679, 2016) some technical modifications as e.g. gentle stirring as proposed by Briassoulis et al. (2020) could be considered to assure the medium is well oxygenated.

Although both tests worked well it is recommended to use the test scenario that is most environmentally relevant for the purpose and to choose well the matrices, especially the sediment used. Sediments of different origin and quality most likely will differ in their microbial activity.

#### 4.1.2 Mesocosm experiments

The testing of the biodegradation of plastic in mesocosms of a larger volume was demonstrated as a viable complement or even alternative to field tests and allowed tests with natural matrices without access to running seawater (Lott et al., 2020). The *t*_0.5_ of the three polymers modelled from disintegration measurements in the open system tests in mesocosms (Figure 5) were about two to four times higher (excluding PBSe, see below) than the ones calculated from the CO_2_ evolution in the lab tests (Figure 3) with the same temperature applied, which confirms the laboratory tests as optimized compared to field tests (even though oxygen availability might have been varying). This fact also confirms our assumption that using the degree of disintegration of samples that are well protected from physical impact as a proxy for biodegradation is well suited to measure the biological processes at the material in the open systems of mesocosms and field experiments rather than a mere physical deterioration of the plastic.

In the mesocosm tests, the variability in the degree of disintegration of single specimens was high even within one tank system, but also between the three replicate tanks and between years, reflected in the sometimes wide 95 % confidence intervals (CI) (Figure 4) as a common feature of all tests. This is attributed to the patchiness of the microbial community, also known from natural sediment environments (e.g. Böer et al., 2009). As can be seen from the shape of the violin plots, the variability within one test is higher in most of the scenarios where degradation was slower, thus half-life higher. For most tests, we applied a set of triplicates for each sampling interval which was just sufficient to obtain a statistically significant basis for modelling. However, this is the absolute minimal replication. Leaving a part of the samples longer deployed in the field tests than planned, led to an insufficient replication, especially considering the high variability observed in all tests. We therefore recommend using 4 or 5 replicates per interval to balance for variability. Furthermore, as seen when comparing the predictions from Asia and the Mediterranean Sea, the estimation of half-life is more precise when higher disintegration (at least about 75%) is reached.

Although observationally different, half-lives were not significantly different between the benthic and eulittoral habitat for the positive control PHA. Also for PBSeT, no consistent significant differences could be found for the half-lives between habitats. Only in the eulittoral test, the half-life in year 2 was significantly lower than in year 1. PBSe showed significantly different half-lives in the benthic and the eulittoral tests. However, very low values for the maximum disintegration in the benthic habitat (1.7% after 237 d) make the prediction of the half-life for this treatment unprecise and the *t*_0.5_ predicted by the model should be taken rather as a rough indication and interpreted with caution. Comparing the performance of the three biodegradable materials in the different mesocosm tests the positive control PHA did disintegrate fastest in the benthic habitat whereas PBSe and PBSeT disintegrated fastest in the eulittoral habitat.

#### 4.1.3 Field tests

The half-lives in the Mediterranean field tests were higher than in the lab for the benthic test for all three polymers, about six times for PHA and about four and eight times higher for PBSe and PBSeT in the eulittoral, respectively. The field test in the Mediterranean Sea revealed significant differences between habitats and, compared with the tests in SE Asia, between climate zones. For all three biodegradable polymers PHA, PBSe, and PBSeT, disintegration was faster in the eulittoral than in the benthic. No significant disintegration was observed in the ultraoligotrophic setting of the Mediterranean pelagic. In the open water of the pelagic, the abundance of microbes is several magnitudes less (e.g. Schmidt et al., 1998) than in seafloor sediments, thus the overall activity of the microbial community is considered much lower. In the tropical waters of SE Asia, disintegration in the benthic was four to fourteen times faster than in the tests in the Mediterranean at the Island of Pianosa.

Marine (and aquatic) biodegradation tests in general often use water as the only matrix (e.g. ASTM D6691, 2017) and give low rates as results (e.g. Bagheri et al., 2017). For biodegradable plastics, we consider this test scenario as the least environmentally relevant given that most biodegradable polymers have a density higher than water and will sink (or float at the water surface if the bulk density of a plastic object (e.g. a closed bottle or foamed material) is <1). However, a water column test is motivated for plastic items typically applied in the open water such as in aquaculture or fisheries.

### 4.2 Temperature differences explain well different biodegradation rates

Temperature is considered one of the most important environmental factors influencing the biodegradation rate. Pischedda et al. (2019) found a significant exponential relation between temperature and the biodegradation rate of a biodegradable plastic material in lab experiments with soil, in accordance with the Arrhenius equation, with the limitation of only three data points and thus no degree of freedom. For a rough estimation of the relationship between biodegradation rate and temperature in our experiments, we therefore applied an exponential model, confirming a significant negative relationship (*F*(1, 8) = 17.654, *p* < 0.003) between the predicted half-life and the average temperature in the mesocosm and field tests (Asia = 28.5 °C, Mediterranean = 17.5 °C, Mesocosm = 20.5 °C). To increase datapoints to a more reasonable number, we did not differentiate between the different polymer types.

### 4.3 Other factors that determine biodegradation

During the same EU project Open-Bio, Briassoulis et al. (2019) conducted similar field tests with the same test materials at a fish farm in Greece in a comparable temperature regime and reported complete disintegration of all samples in the benthic tests after 9 months, roughly accounting for a half-life of 140 d (estimated from the graphs provided) as compared to 738 d (PHA), 908 d (PBSe) and 1070 d (PBSeT) in our Mediterranean benthic tests. This indicates that the trophic level of the habitat might also have a positive influence on the biodegradation activity (and rate). In our study, most of the nutrient-related parameters (*sensu* Weber et al., 2012) such as concentrations of N and P compounds were low in the three different settings. Manipulative tank experiments could be used to investigate the relationship between nutrient concentration and biodegradation rate.

The abundance of microbes and their community composition, especially in tests involving sediments, might differ strongly, e.g. dependent on factors such as grain size (Ahmerkamp et al., 2020). Different sediment types such as e.g. coarse permeable sand, fine silt, or mud due to their different permeabilities might favor strongly differing metabolic pathways, e.g. depending on the presence or absence of oxygen as an electron acceptor. Some polymers might also behave differently under oxic or anoxic conditions than others. As reported recently (Lott et al., 2020), we also observed a different disintegration performance within the same sample exposed in the HYDRA® frames (SI Figure 1). Due to the tightly adhering frame, the margin of the exposed plastic film (which was not accounted for in the disintegration measurements) supposedly had less exchange with the surrounding water, and the microbes there presumably experienced hypoxic or anoxic conditions.

PHA and PBSe showed a contrasting trend in the performance in the benthic and eulittoral tests in the mesocosms experiment: PHA was disintegrating faster in the benthic than in the eulittoral and PBSe faster in the eulittoral than in the benthic. In the field tests, PHA and PBSeT disintegrated faster in the eulittoral than in the benthic, also the half-life predictions for PBSe followed this trend, but differences were not significant. In line with the results of the mesocosm tests discussed above, this is an indication that the comparison of a test material with a reference material has to be done with some caution and the application of a material as a positive control is critically scrutinized. Some well-degradable materials might perform differently under certain conditions than expected, making the desired intercalibration of test results e.g. between habitats or climate zones based on one material difficult and conclusions should be drawn with caution.

### 4.4 Creating comparable data for LC(I)A

The predictive modelling of half-lives we present here gives the opportunity to assign a numerical value for the biodegradation performance in a defined scenario (e.g. Mediterranean Sea/Pianosa Island/benthic sand) as a material property. Data provided in this manner by us and studies to come should be used for the compilation of a catalogue of biodegradable materials and their properties with regard to their environmental performance that can be used for statistical comparisons of different materials in the same habitat, or one material in different environmental scenarios. These specific half-lives will also be suited to enable further mathematical modelling e.g. for environmental risk assessment and the life cycle (impact) assessment of products. Furthermore, such a catalogue could help industry, public administration, and NGOs to base their decisions for or against certain materials and their application on facts.

The principle of half-life, however, is difficult to communicate and to perceive to non-specialists e.g. the general public or policy makers. The question ‘How long does it take?’ (until a certain object is completely degraded) cannot be readily answered by the specific half-life. Dilkes-Hoffman et al. (2019) compared the results of PHA biodegradation tests under marine conditions in a meta-analysis from eight former studies. To achieve comparability, they recalculated the data to a rate of mass loss over time per exposed surface area (Equation 3) based on the assumption that in biodegradable solids the processes causing the degradation are only happening at the surface of the object:

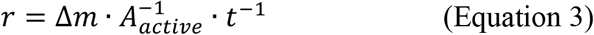

where *m* is mass, *A* the surface area exposed, and *t* the time.

From these data, they also estimated the lifetime, i.e. the time needed for an *object* to completely degrade. The 95 % CI of the biodegradation rates derived from all studies considered was 0.04 - 0.09 mg cm^−2^ d^−1^. We applied their calculation method to our experimental results on PHA, using the volume of the films (area × thickness) and the specific gravity *ρ* of the material (*ρ*_PHA_ = 1.3). We obtained rates within or close to this range only for our fastest scenarios, PHA exposed in the benthic test in SE Asia (0.051 mg cm^−2^ d^−1^), and in lab tests (0.012 mg cm^−2^ d^−1^ for eulittoral and 0.024 mg cm^−2^ d^−1^ for benthic tests). For the other test scenarios, the biodegradation rates were one order of magnitude lower (SI Table 9).

If we apply the minimum and maximum biodegradation rates derived from the results of our beach and seafloor field tests in the Mediterranean Sea and in tropical SE Asia (0.051 - 0.004 mg cm^−2^ d^−1^; SI Table 9) to estimate of the lifetime of the PHA objects as in figure 3 of Dilkes-Hoffman et al. (2019) it results in lifetimes 1.8 – 10 times higher, ranging from 2 – 20 months for a 35 μm thick shopping bag to 2.7 - 36 years for a PHA bottle with 800 μm thickness to 4 – 54 years for PHA cutlery (~1300 μm). It has however to be taken into account, that the calculation of the rate by Dilkes-Hoffman et al. (2019) is based on the initial surface of the object which in reality will change during the biodegradation process. Very likely, the surface roughness will increase and the surface-to-volume ratio will become higher and thus the gross degradation of the object or its fragments will accelerate, as was also shown by Chinaglia et al. (2018). Therefore, these estimations should be taken as conservative for the environments considered. On the other hand, given the strong temperature dependence of the biodegradation rate, it can be assumed that in colder environments as the deep sea or in polar regions the lifetime will be higher.

The dependence of the (bulk) biodegradation rate on the surface-to-volume (or mass) ratio was mentioned by Modelli et al. (1999) who, in a study with another focus, compared powder (grain size 1 μm) and film of PHA (4 × 4 × 0.025 cm^3^) in a soil biodegradation test according to ASTM 5988 (2003) and interpreted the rates as different. The authors missed to numerically address the surface-to-volume ratio in this comparison and stated that the initial rate (11 % of the 0.5 g bulk polymer in 1 d) was 90 times higher for powder deduced from the slope of the linear fits. However, if their data is re-calculated with Equation 3, the rates of 0.023 [0.056 g · d^−1^; A = 2400 cm^2^] for powder (simplified assuming spherical particles) and 0.020 [0.132 g · 210 d^−1^; A = 32 cm^2^] for film are almost identical and the difference in the bulk rates (90 times) is well explained by the difference in surface area (75 times).

Chinaglia et al. (2018) tested also in soil (ASTM D5988, 2003) lab experiments at 28 ±2 °C, PBSe powder of different grain sizes and found the maximum rate for 1 g of sample at 97 mg C_polymer_ d^−1^ (and the total surface in dependence of grain size where half the maximum rate was reached at 1122 cm^2^). If equation 3 is applied to their data, the areal rates for PBSe particles (0.05 – 0.18 mg · cm^2^ · d^−1^) are about 10 to 600 times higher than our results for PBSe film (25 μm thickness) under marine conditions (SI Table 10).

Taking into account that biodegradation of a solid object is occurring at the surface the specific half-lives modelled from results from our tests with film can be converted to erosion rates in μm per year *yr* by dividing half the film thickness *h* by the half-life *t*_0.5_:

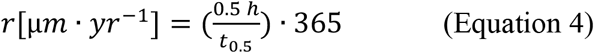

These values are surface-independent and density-independent and can be applied to three-dimensional objects with parallel surfaces as e.g. most packaging such as bottles etc. with the only object-specific parameter to know being the material thickness.

Applying Equation 4, microbial surface erosion (‘micro-bioerosion’) rates derived from our experimental results range from 12.4 – 285.7 μm · yr^−1^ for PHA, 5.7 – 31.1 μm · yr^−1^ for PBSeT and 1.7 – 46.0 μm · yr^−1^ for PBSeT (see Table 4).

The micro-bioerosion rate in the respective habitat divided by the wall thickness in μm of an object will provide estimated lifetimes that can be used for environmental risk and life cycle assessments. Further experiments on solid objects rather than the film will be useful to validate these estimations.

The numbers presented here might also help to clarify the assumption that degradation of biodegradable plastic in the marine environment is occurring ‘much more slowly’ and give useful input to the statement that ‘the degree to which “biodegradable” plastics actually biodegrade in the natural environment is subject to intense debate’ (UNEP, 2015). Our data and also previous studies show that there are biodegradable plastic materials that do degrade relatively fast in the natural marine environment given their functional properties in the applications they were designed for. A stable, durable, and resistant plastic item that performs well during use is unlikely to ‘disappear’ within a few days or weeks once lost or littered to the natural environment. This also underpins the urgency to apply all possible means to prevent any plastic material from entering the natural environment, being it biodegradable or not. Plastic lost to the environment is pollution, even if biodegradable. However, biodegradable plastics are less likely to accumulate or persist than conventional plastics.

## Supporting information

Supplemental Information to LOTT et al

## 5 Acknowledgements

Deep thanks to the student assistants and interns of HYDRA for their help during experiment preparation and sampling. Special thanks for technical work to Eskil Salis Gross for sediment characterization, to Nora Pauli, Alexandra Belitz and Esther Thomsen for image analysis. We thank the National Park Tuscan Archipelago, Portoferraio for granting access to the protected area of the Island of Pianosa with the research permit n.3063/19.05.2014 and following. Dott. Emiliano Somigli and his staff are gratefully acknowledged for their support and for granting access to Terme San Giovanni to perform the eulittoral tests. We thank Giorgio Vendetti from Hotel Mirage, Marina di Campo, for providing meteorological data (www.elbaexplorer.com). The Government of the Republic of Indonesia, Ministry of Research, Technology and Higher Education, RISTEK-DIKTI, Jakarta is gratefully thanked for granting the research permits no. 71 and 72/SIP/FRP/E5/Dit.KI/III/2017 and extensions to C.L. and M.W.. C.L. and M.W. express their thanks to University Sam Ratulangi, Manado welcoming them as guest researchers. Thanks to Ilaria Reggi, Anna Clerici, Marco Perin, and staff of Coral Eye Resort and Coral Research Outpost, Bangka Island, Sulawesi Utara, Indonesia for technical support and maintenance. The study in the Mediterranean Sea was conducted within the European Union’s FP7 project Open-Bio and has received partial funding under the grant agreement no KBBE/FP7EN/613677 and by BASF SA, Ludwigshafen, Germany. Thanks also to Novamont S.p.A., Novara, Italy for providing film for tests during the Open-Bio project. We also thank our partners of the WP5 of the Open-Bio project for valuable discussions on the outcomes of the tests.

