## Supplemental Information to LOTT et al for "Half-life of biodegradable plastics in the marine environment depends on material, habitat, and climate zone"

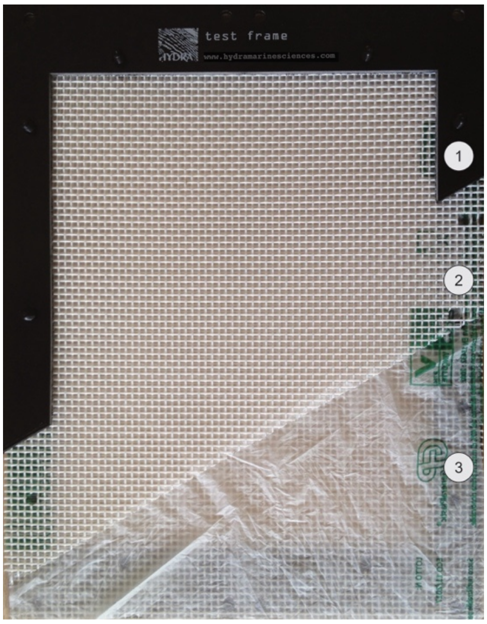

**SI Figure 1. HYDRA test frame exemplarily cut open for demonstration:** (3) Test material as film, covered by (2) mesh (PET) and held by (1) plastic (PE) frame. (modified from Lott et al., 2020)

SI Table 1. **Mesocosm experiments with Mediterranean matrices:** Sampling dates of all samples in the three habitats eulittoral (intertidal beach scenario), pelagic (water column scenario), and benthic (sublittoral; seafloor scenario) in the mesocosm test of the experiment of year 1 and year 2. * The last sampling interval of the second-year experiment was only 1.5 months due to the termination of the project.

| Eulittoral | | | | | | | |
|---|---|---|---|---|---|---|---|
| year 1 | | | | year 2 | | | |
| interval | date | months | days | interval | date | months | days |
| $t_0$ | 18.04.2014 | 0 | 0 | $t_0$ | 28.04.2015 | 0 | 0 |
| $t_1$ | 10.07.2014 | 2.5 | 83 | $t_1$ | 15.07.2015 | 2.5 | 78 |
| $t_2$ | 16.09.2014 | 5 | 151 | $t_2$ | 30.09.2015 | 5 | 155 |
| $t_3$ | 05.12.2014 | 7.5 | 231 | $t_3$ | 18.12.2015 | 7.5 | 234 |
| $t_4$ | 20.02.2015 | 10 | 308 | $t_4$ | 01.03.2016 | 10 | 308 |
| $t_5$* | 04.03.2016 | 22 | 686 | | | | |
| Pelagic & Benthic | | | | | | | |
| year 1 | | | | year 2 | | | |
| interval | date | months | days | interval | date | months | days |
| $t_0$ | 28.07.2014 | 0 | 0 | $t_0$ | 04.07.2015 | 0 | 0 |
| $t_1$ | 07.10.2014 | 2.5 | 71 | $t_1$ | 27.09.2015 | 2.5 | 85 |
| $t_2$ | 22.12.2014 | 5 | 147 | $t_2$ | 16.12.2015 | 5 | 165 |
| $t_3$ | 13.03.2015 | 7.5 | 228 | $t_3$ | 02.03.2016 | 7.5 | 242 |
| $t_4$ | 06.06.2015 | 10 | 313 | $t_4$ | 18.05.2016 | 10 | 319 |
| $t_5$* | 05.06.2016 | 22 | 678 | | | | |

**SI Table 2. Mediterranean field experiments:** Sampling dates of all samples in the three habitats eulittoral (intertidal beach scenario), pelagic (water column scenario), and benthic (sublittoral; seafloor scenario) in the field test of the experiment of year 1 and year 2. *In order to learn more about the disintegration process over a longer exposure time, the last (additional) sampling interval of the first-year experiment was 22 months.

| Eulittoral | | | | | | | |
|---|---|---|---|---|---|---|---|
| year 1 | | | | year 2 | | | |
| interval | date | months | days | interval | date | months | days |
| $t_0$ | 22.09.2014 | 0 | 0 | $t_0$ | 29.09.2015 | 0 | 0 |
| $t_1$ | 10.12.2014 | 2.5 | 79 | $t_1$ | 15.12.2015 | 2.5 | 77 |
| $t_2$ | 22.02.2015 | 5 | 153 | $t_2$ | 27.02.2016 | 5 | 152 |
| $t_3$ | 08.05.2015 | 7.5 | 228 | $t_3$ | 23.05.2016 | 7.5 | 238 |
| $t_4$ | 27.07.2015 | 10 | 308 | $t_4$* | 26.06.2016 | 9* | 270 |
| Pelagic & Benthic | | | | | | | |
| year 1 | | | | year 2 | | | |
| interval | | months | days | interval | | months | days |
| $t_0$ | 22.09.2014 | 0 | 0 | $t_0$ | 29.09.2015 | 0 | 0 |
| $t_1$ | 09.12.2014 | 2.5 | 78 | $t_1$ | 15.12.2015 | 2.5 | 77 |
| $t_2$ | 23.02.2015 | 5 | 154 | $t_2$ | 27.02.2016 | 5 | 151 |
| $t_3$ | 09.05.2015 | 7.5 | 229 | $t_3$ | 23.05.2016 | 7.5 | 237 |
| $t_4$ | 27.07.2015 | 10 | 308 | $t_4$* | 26.06.2016 | 9* | 271 |

**SI Table 3:** Formulas used to calculate half-life with the estimated model parameters *a* (*y*-intercept) and *b* (slope) for different link-functions

| Link-function | Formula |
|---|---|
| logit | $t_{0.5} = -a / b$ |
| cloglog | $t_{0.5} = (\log(-\log(0.5)) - a) / b$ |
| cauchit | $t_{0.5} = -a / b$ |
| loglog | $t_{0.5} = -(\log(-\log(0.5)) - a) / b$ |

**SI Table 4:** Empirical *p*-values for comparisons of the half-life for polymer PHA in the mesocosm experiment between all groups. *p*-values were adjusted for multiple comparisons with the Holm method. Significant *p*-values are bold.

| **Group 1** | **Group 2** | ***p*-value** |
|---|---|---|
| Benthic | Eulittoral | **0.0180** |
| Benthic | Pelagic | **0.0024** |
| Eulittoral | Pelagic | 0.1160 |

**SI Table 5:** Empirical *p*-values for comparisons of the half-life for polymer PBSeT in the mesocosm experiment between the two years within habitats. *p*-values were adjusted for multiple comparisons with the Holm method. Significant *p*-values are bold.

| **Group 1** | **Group 2** | ***p*-value** |
|---|---|---|
| Benthic year 1 | Benthic year 2 | 0.0555 |
| Benthic year 1 | Eulittoral year 1 | 0.2155 |
| Benthic year 1 | Eulittoral year 2 | **0.0081** |
| Benthic year 1 | Pelagic year 1 | 1.0000 |
| Benthic year 1 | Pelagic year 2 | **0.0081** |
| Benthic year 2 | Eulittoral year 1 | 1.0000 |
| Benthic year 2 | Eulittoral year 2 | **0.0001** |
| Benthic year 2 | Pelagic year 1 | 1.0000 |
| Benthic year 2 | Pelagic year 2 | **0.0001** |
| Eulittoral year 1 | Eulittoral year 2 | **0.0071** |
| Eulittoral year 1 | Pelagic year 1 | 1.0000 |
| Eulittoral year 1 | Pelagic year 2 | **0.0376** |
| Eulittoral year 2 | Pelagic year 1 | 0.0707 |
| Eulittoral year 2 | Pelagic year 2 | 0.0532 |
| Pelagic year 1 | Pelagic year 2 | 0.0707 |

**SI Table 6:** Empirical *p*-values for comparisons of the half-life for polymer PHA in the field experiments between all groups. *p*-values were adjusted for multiple comparisons with the Holm method. Significant *p*-values are bold.

| **Group 1** | **Group 2** | ***p*-value** |
|---|---|---|
| Asia benthic | Med.Sea benthic | $<$**0.0001** |
| Asia benthic | Med. Sea eulittoral | $<$**0.0001** |
| Med. Sea benthic | Med. Sea eulittoral | **0.0161** |

**SI Table 7:** Empirical *p*-values for comparisons of the half-life for polymer PBSeT in the field experiments between all groups. *p*-values were adjusted for multiple comparisons with the Holm method. Significant *p*-values are bold.

| Group 1 | Group 2 | *p*-value |
|---|---|---|
| Asia benthic | Med. Sea benthic | **<0.0001** |
| Asia benthic | Med. Sea eulittoral | **<0.0001** |
| Med. Sea benthic | Med. Sea eulittoral | **0.0008** |

**SI Table 8:** Empirical *p*-values for comparisons of the half-life for polymer PBSe in the field experiments between all groups. *p*-values were adjusted for multiple comparisons with the Holm method. Significant *p*-values are bold.

| Group 1 | Group 2 | *p*-value |
|---|---|---|
| Asia benthic | Med. Sea benthic | **0.0024** |
| Asia benthic | Med. Sea eulittoral | **0.0465** |
| Med. Sea benthic | Med. Sea eulittoral | 0.7233 |

**SI Table 9: Areal biodegradation rates and lifetimes (*sensu* Dilkes-Hoffmann, 2019) of PHA films (85 µm) under different marine conditions.** LAB = laboratory tests; MESO = mesocosm tests; FIELD = field tests; MED = Mediterranean Sea; ASIA = tropical SE Asia; EUL = eulittoral (beach scenario); BEN = benthic (sublittoral sand bottom scenario); PEL = pelagic (open water scenario); [a] = film of 8 cm$^2$; [b] = film of 3.1 cm$^2$; [c] = film of 320 cm$^2$.

| | | | **PHA** [85 µm thickness] | |
|---|---|---|---|---|
| **test** | **climate** | **habitat** | **areal rate [mg d$^{-1}$ cm$^{-2}$]** | **lifetime [d]** |
| **LAB** | | **EUL** | **0.012472** | **443**[a] |
| | | **BEN** | **0.023835** | **232**[b] |
| **MESO** | **MED** | **EUL** | **0.003750** | **1473**[c] |
| | | **BEN** | **0.007742** | **714**[c] |
| | | **PEL** | **0.002216** | **2493**[c] |
| **FIELD** | **MED** | **EUL** | **0.006623** | **834**[c] |
| | | **BEN** | **0.004226** | **1307**[c] |
| | **ASIA** | **BEN** | **0.050875** | **109**[c] |

**SI Table 10: Areal biodegradation rates [mg d$^{-1}$ cm$^{-2}$] (*sensu* Dilkes-Hoffmann, 2019) of PHA films (85 µm), PBSe films (25 µm) and PBSeT films (25 µm) under different marine conditions.** Eulittoral = beach scenario; Benthic = sublittoral sand bottom scenario; Pelagic = open water scenario.

| **Polymer** | **Habitat** | **Year** | **Lab** | **Mesocosm** | **Mediterranean Sea** | **SE Asia** |
|---|---|---|---|---|---|---|
| PHA | Benthic | | 0.023835 [0.024776, 0.022983] | 0.007742 [0.009962, 0.005173] | 0.004226 [0.004360, 0.004116] | 0.050875 [0.056493, 0.046273] |
| PHA | Eulittoral | | 0.012472 [0.014225, 0.009998] | 0.003750 [0.005143, 0.002093] | 0.006623 [0.008311, 0.004729] | |
| PHA | Pelagic | | | 0.002216 [0.003220, 0.001069] | | |
| PBSeT | Benthic | 1 | | 0.001005 [0.001810, 0.000588] | | |
| PBSeT | Benthic | 2 | | 0.001753 [0.001996, 0.000641] | | |
| PBSeT | Benthic | | 0.003843 [0.004424, 0.003100] | | 0.000980 [0.001050, 0.000893] | 0.003893 [0.004257, 0.003614] |
| PBSeT | Eulittoral | 1 | | 0.002141 [0.004384, 0.001301] | | |
| PBSeT | Eulittoral | 2 | | 0.005322 [0.006451, 0.002223] | | |
| PBSeT | Eulittoral | | | | 0.001110 [0.001136, 0.001079] | |
| PBSeT | Pelagic | 1 | | 0.001469 [0.002941, 0.000716] | | |
| PBSeT | Pelagic | 2 | | 0.003491 [0.003763, 0.002688] | | |
| PBSe | Benthic | | 0.004173 [0.004248, 0.004101] | 0.000299 [0.000373, 0.000224] | 0.000553 [0.000817, 0.000250] | 0.007876 [0.008401, 0.007315] |
| PBSe | Eulittoral | | | 0.005272 [0.005878, 0.004718] | 0.000646 [0.001139, 0.000151] | |
